# Adult regenerative defects arise from discordant scaling of signal dependent growth and patterning

**DOI:** 10.64898/2026.04.28.721472

**Authors:** Pietro Tardivo, Fabrizio Olmeda, Martin Nuamah Asare, Jessica Lange, Dunja Knapp, Andre Fischer, Yuka Taniguchi-Sugiura, Prayag Murawala, Edouard Hannezo, Elly M. Tanaka

## Abstract

Orders of magnitude distinguish an organ’s size in adulthood from its size when patterning was established during embryonic development. The prospect of engineering adult organ regeneration raises the fundamental question of whether regenerative stem cell patterning should be elicited at embryonic or adult scale. Axolotls regenerate their limbs at all stages of post-embryonic life, encompassing an order of magnitude increase in animal size, but the mechanisms allowing this robust capacity remain unclear. Limb regeneration occurs by the formation of an embryonic-like progenitor zone, the blastema, whose dimensions increase with animal size, suggesting that at least some aspects of limb regeneration must scale. Here, by combining spatial transcriptomics, biophysical modelling, quantitative imaging, and functional perturbations, we found distinct scaling signatures among key signaling pathways, arguing against a body-size-dependent hormonal scaling mechanism. SHH signaling showed partial scaling that saturated in largest adult sizes, with a correspondingly early termination of blastema growth while Wnt9a-modulated digit patterning scaled at all animal sizes. As a result of this differential scaling, digit patterning wavelength was mismatched with respect to domain growth in the largest blastemas, preventing the addition of the last digit. In agreement with this model, regenerative failure can be rescued by timed supplementation of SHH. Our results show that coordinated scaling of morphogen signaling is a key requirement for adult organ regeneration.

## Main

Axolotl limb regeneration is a post-embryonic process in which limb amputation stimulates formation of a progenitor pool called blastema that recapitulates the limb development gene expression network ^1,2^. This developmental network relies on long-range signaling from multiple, localized signaling centers including a posterior SHH and an anterior FGF8 expression domain that form a positive feedback loop for limb bud growth ^2,3^. Regeneration can occur in animals of different ages and sizes from 2.5 cm to 25 cm long animals ^4,5^, which encompasses larval, juvenile and adult stages. Importantly, regeneration does not involve the formation and patterning of an embryonically-sized blastema. Rather, previous work showed that the width of the mid-bud stage blastema ("limb field") scales allometrically with animal length ^2,5^, representing a >6-fold expansion of the limb field, which suggests that scaling mechanisms must exist to adapt regeneration to different animal and tissue sizes. Colorimetric detection of *Shh* (source) and *Ptch1* (target gene) mRNAs suggested that the *Shh* expression domain expands linearly with limb-field size ^4,5^, without a corresponding increase in *Ptch1* expression length scale. This leads to a model where the size of the morphogen production domain, rather than its signaling range, is a key feature of blastema scaling. Other modeling work proposed body-size dependent, static scaling of SHH and FGF8 morphogen gradients as a mechanism to account for scaling of blastemas ^5^. Beyond these outstanding questions on the mechanisms and nature of SHH scaling, accurate limb regeneration requires scaling of multiple other developmental events, such as the scaling of digit periodicity with tissue size, yet neither digit patterning nor regeneration fidelity have been studied across animals of different size. Overall, whether and how regenerative scaling of the limb signaling network occurs remains unsolved.

Previous studies of developing tissues have examined scaling of several morphogen pathways ^6–10^. Studies of *Drosophila* wing development have shown that Dpp signaling scales with tissue size during wing growth ^11–13^, but Hedgehog signalling does not ^11^. Comparison of the chicken and zebra finch neural tube showed that cell-intrinsic differences in signal-sensitivity mediate scaling of SHH signaling across the two species ^14^. Within a single species, zebrafish embryo perturbations demonstrated that size-dependent feedbacks can rescale SHH and Nodal signaling after surgical size-reductions ^8,15^. In these developmental contexts, morphogen scaling is fundamental for ensuring robust patterning despite natural variation and tissue growth ^16^. Regeneration, however, offers perhaps the most remarkable opportunity to study scaling as, within the same genetic background, regenerating tissues must rescale signaling to dimensions that far exceed those of development. For example, in the regenerating zebrafish fin, scaled expression of Fgf ligands, coupled to proliferation-dependent advection, has been proposed to control regenerative growth across millimeters. Planarians can reestablish head-to-tail body patterning in adult sizes that are an order of magnitude larger than during development. Yet, this arises due to rescaling of a pre-existing Wnt morphogen gradient ^17,18^. In contrast, in the axolotl limb, signaling gradients are absent before amputation and must be established *de novo* in the blastema. This provides a unique context to study formation and scaling of morphogen signaling and patterning at post-embryonic scales.

Here, we obtained a global view of signal pathway scaling during axolotl limb regeneration using spatial transcriptomics of limb blastemas after lower arm amputation in different sized animals. We identified clear but distinct scaling signatures across multiple signaling pathways and found that scaling involves not only an increase in the size of morphogen production domains, but of their target domains, which signifies expanded signaling range. Using high resolution, quantitative spatial profiling of *Shh* mRNA and its receptor and target gene *Ptch1,* we find a linear 4-fold expansion in the SHH signaling range in ∼2 to 8 cm animals. Interestingly, SHH signaling scaling plateaus in animals larger than 10 cm animals, which can be quantitatively described by an expansion-repression mechanism. We also show that this loss of SHH scaling has functional consequences, with larger animals displaying reduced blastema growth and abnormally low digit counts. In contrast to under-scaling of blastema growth and SHH signalling, we find that digit specification, which occurs via a periodic patterning mechanism, scales at all tissue sizes. However, digit number depends not only on patterning wavelength but also on the continued growth of the blastema. Indeed, digit loss in large tissues can be explained by insufficient field growth relative to a correctly scaled digit patterning wavelength. Accordingly, we could rescue digit defects either by timely supplementing SHH or via an upper arm amputation, which yields sustained blastema growth. These results highlight the importance of coordinating morphogen signal scaling in regenerative tissues to obtain correctly patterned body structures. Altogether, we provide a roadmap to rescue regenerative failures due to discoordinated signal scaling.

### Spatial sequencing identifies scaling features in limb regeneration

Axolotls are capable of regenerating limbs over a broad range of sizes ^4,5,19^, raising the question of whether and how gene expression during regeneration scales with tissue size. To examine this question, we performed lower arm amputations in three different axolotl sizes and carried out spatial transcriptomic sequencing of regenerating blastemas (see *Methods*, Fig. 1a). We analyzed eight batches, comprising five to six cross-sections from animals of different sizes, and covering three regeneration stages. As our focus centers on the mesenchyme where many key signaling activities occur ^1,2^ (Fig. S1a), we excluded epithelial populations from further analysis (Fig. S1a, S1b, *Methods*).

**Figure 1:**
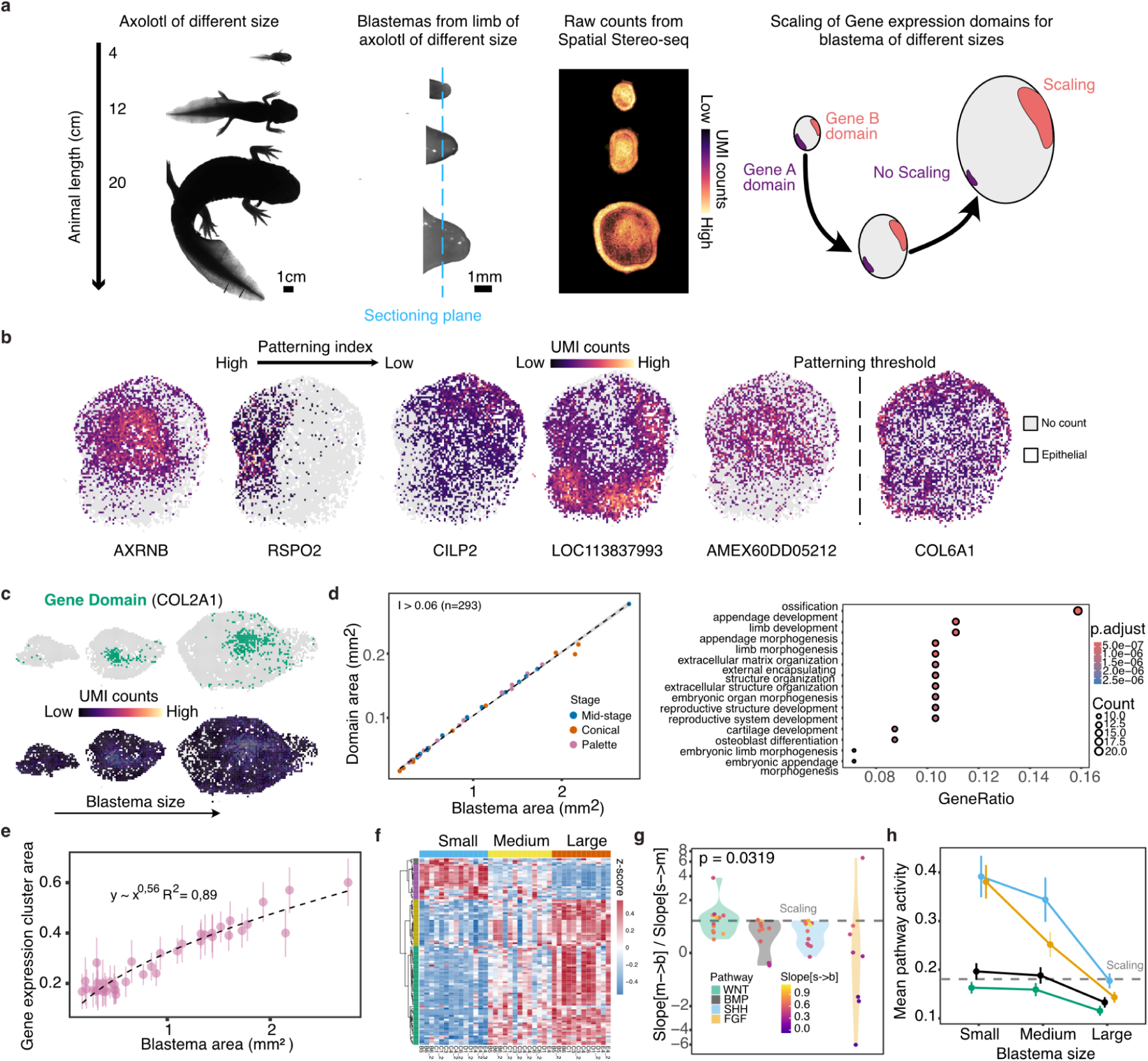
**a.** Left: representative images of 4,12 and 20cm long axolotls from which forelimbs were amputated to generate blastemas for Stereo-seq. Middle left: forelimb blastemas from different-sized animals; dashed line schematizes sectioning orientation. Middle right: heatmap of total UMI counts in blastema sections derived from the blastemas which are the basis of further analysis. Right: schematized version of different possibility of scaling of gene expression domain: Domain can scale proportionally to the blastema area or fail to scale. **b.** Spatial distribution of UMI counts of six different genes selected and ordered by their patterning index (*Methods*) from one blastema profiled with Stereo-seq (bigger blastema in the B6 batch). Each dot corresponds to a bin in the Stereo-seq data (size of 25 μm²). The patterning threshold sets the limit below which gene domains are disregarded from the analysis in **d**. **c.** The typical gene domain from the C3 batch detected from UMI counts for increasing larger blastemas in the same batch (*Methods*). UMI counts are normalized within the batch. **d,** Left: Gene domain size of the genes which are mostly patterned (293 in this plot) for increasingly larger blastemas. Different thresholds are shown in Fig. S1c. Each dot corresponds to the average domain for all the genes with patterning index greater than the threshold in one blastema. Right: GO enrichment analysis performed on the most patterned genes reveals that patterned genes overlap with limb development and regeneration programs. **e:** Scaling of the normalized average cluster size for all the clusters identified with clustering algorithms for the mesenchyme (*Methods*). Individual dots correspond to the average size of clusters identified in *Methods* for each blastema. We select only clusters which are present in at least one small, medium and large blastema. **f.** Differential expression analysis and module identification for the highly variable genes between smaller and large size blastemas. GO terms of downregulated genes in larger blastemas are shown in Fig. S1E. Gene names are provided in Extended Data Table 1. **g.** Violin plot of the ratio between the coefficient from linear regression between the average domain of individual genes from small to medium sized blastemas and medium to large sized blastemas. A value of one of the ratios corresponds to perfectly linear scaling. Individual gene scaling is given in Fig. S1g. Positive and negative values correspond to genes of which the domain overscale or underscale with the blastema area. The color of the dot is the coefficient of linear regression between small to large blastemas. The dashed line (Scaling) refers to the ratio between slopes being equal to one. **h.** Mean pathway activity: cell-averaged pathway expression per blastema, further averaged across blastemas within each size class. The dashed line (Scaling) is a reference for linear scaling of the mean activity with respect to the blastema size.

We first asked how expression of genes with non-uniform spatial profiles scale across blastemas of different sizes. We categorised genes with respect to their spatial pattern by assigning them a patterning index (Fig. 1b, *Methods*). Genes with high patterning index typically have well-defined spatial domains of gene expression that are suitable for a scaling analysis (Fig. 1c). After identifying the highly variable genes (HVGs) across all the blastemas (*Methods*), we selected those with the highest patterning index and computed the average domain area for each blastema section (*Methods*). We found that the average gene domain area is a linear function of the area of the blastema (Fig. 1d). This linear dependence is consistent for different choices of the patterning index threshold (Fig. S1c). Gene Ontology (GO) analysis of these non-uniform genes revealed that they are preferentially associated with limb development and skeletal differentiation (Fig. 1d, right panel).

As an alternative way of analyzing scaling in our dataset considering both uniform and non-uniform genes, we clustered bins as a function of transcriptional similarity (Fig. S1d, *Methods*), without reference to their spatial position. We then quantified cluster size, i.e. the number of bins belonging to each cluster, as a function of blastema size. Interestingly, we found a sub-scaling behaviour with blastema size, when averaging across clusters (*Methods*), with an exponent of 0.56 (Fig. 1e). Analysis at the level of individual clusters revealed that the majority of clusters behaved similarly to the average, with sublinear scaling of cluster size with blastema area (Fig. 1e, Fig. S1d).

Given the discrepancy in scaling behaviour between HVG-pattern domains versus transcriptomically similar clusters, we asked whether transcripts associated with subscaling were enriched in particular gene functions. To identify candidate transcripts associated with sublinear scaling in an unbiased manner, we performed a differential expression analysis between blastemas of different sizes (this analysis would also identify transcripts whose expression intensity decreased in large animals). We found one group of coexpressed genes whose expression levels are downregulated in larger blastemas, in contrast to most genes which are upregulated (Fig. 1f). Interestingly, GO analysis revealed that the top terms for this group were related to limb development and morphogenesis (Fig. S1e). Given these categories of gene enrichment, we specifically evaluated spatial scaling of WNT, BMP, FGF and SHH signalling pathway members (Fig. 1g, Fig. S1g), which have been implicated in limb regeneration ^2,20,21^. While the spatial domains of transcripts associated with WNT and BMP pathways scaled nearly linearly (slope ratio ∼ 1), those associated with SHH and FGF pathways displayed sublinear scaling (slope ratio < 1), meaning that larger animals tend to show weaker increase of domain size with blastema size (Fig. 1h). Altogether, this analysis revealed that scaling is not a global property of limb regeneration, but rather that different pathways show different scaling behavior.

### SHH-signaling scales via a size-dependent feedback mechanism

Given that SHH is a central regulator of limb regeneration ^2,22^, and is suggested to scale in a sublinear manner by our analysis, we next chose to specifically scrutinize the gene expression features of the SHH pathway (Fig. 2a and Fig. S2a,b). The SHH target genes *Ptch1* and *Ptch2* showed high expression posteriorly that decreased along the posterior-to-anterior axis, while *Gli3* transcripts were distributed throughout the blastemas with a peak in the medial-anterior region. The graded expression of *Ptch* transcripts was consistent with concentration-dependent signaling of SHH ligands from posterior to anterior. We therefore plotted their expression profiles along a posterior-to-anterior axis normalized to blastema size, which showed scaling in smaller sizes but subscaling in large blastemas.

**Figure 2:**
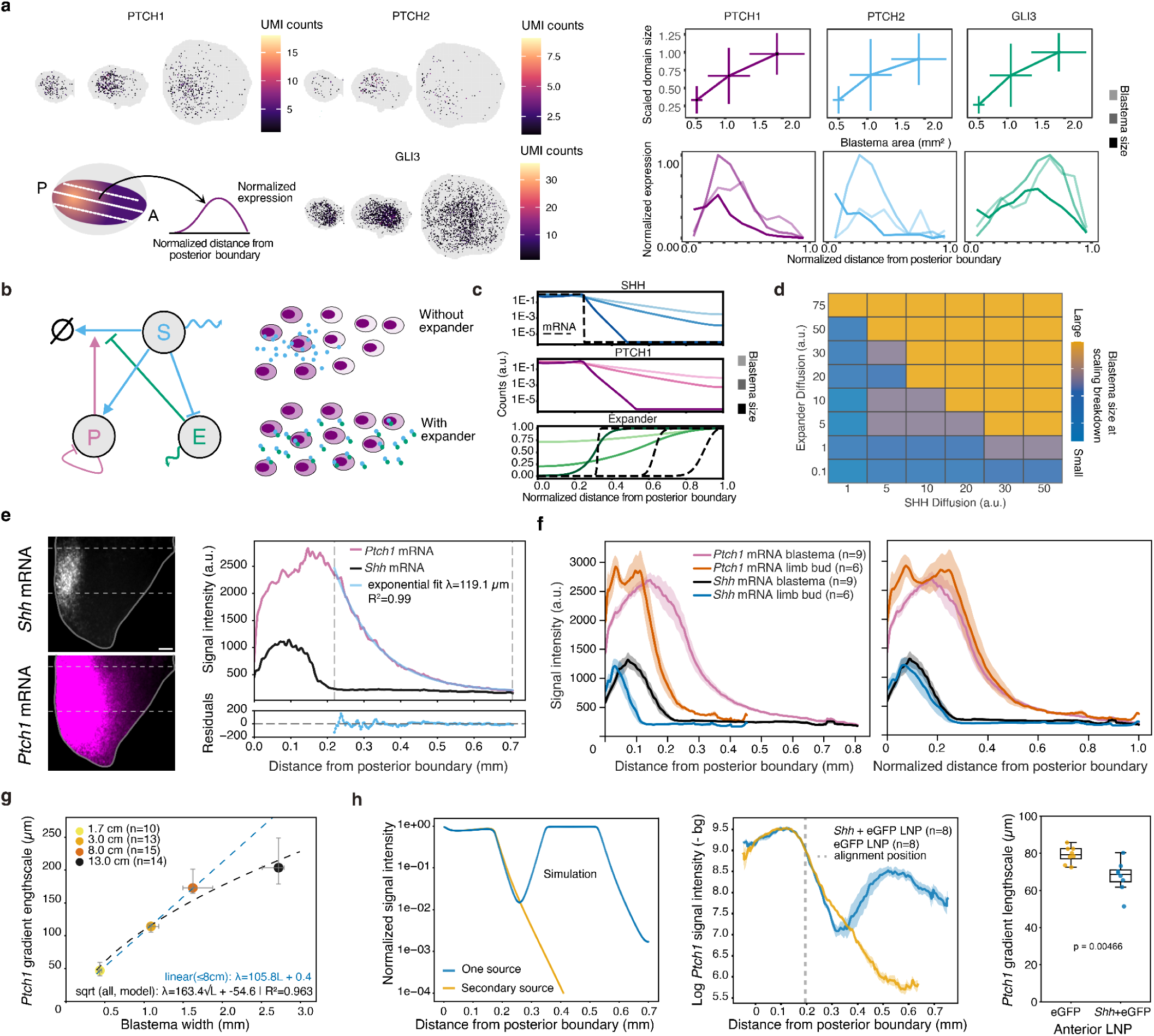
**a:** Spatial expression of selected SHH pathway genes during limb regeneration (left), normalized domain size as a function of blastema area (top right), and expression profiles plotted against normalized tissue width for different blastema sizes (bottom right). Domain size increases with blastema area and reaches a plateau in larger blastemas. **b:** Minimal gene networks consistent with the data between *Shh* (*S*), *Ptch1* (*P*) and the expander (*E*) (*Supplementary Theory*). **c:** Numerical simulations of the model in (**b**) reveals scaling profiles of *Shh*, *Ptch1* and expander. The profile for *Ptch1* mRNA is proportional to the protein up to a scaling factor (*Supplementary theory*). Simulated blastema sizes are L = 100, 200, and 1000 (in simulation units). Expression profiles are normalized to their maximum value. *Shh* and *Ptch1* gradients are shown on logarithmic scales to highlight their exponential decay. **d:** The minimal model predicts a change in length-scale of the gradient and a shift of the *Ptch1* peaks compared to the end of the SHH domain. We show the normalized length at which the length-scale of PTCH1 fails to scale for several values of SHH and expander diffusion. All the other simulation parameters are given in Table 1 of the *Supplementary Theory*. **e:** Extraction of one-dimensional *Ptch1* gradient lengthscales from in situ hybridization images. Left: Representative average projection image of *Shh* and *Ptch1* mRNA expression (*Methods*); dashed lines delimit the band from which signal is extracted. Right: *Shh* and *Ptch1* mRNA signal intensity profiles as a function of distance from the posterior boundary. An exponential decay function is fitted to the *Ptch1* profile (partially constrained background fit, see *Methods*), yielding the characteristic lengthscale λ; residuals of the fit are shown below. **f:** *Ptch1* and *Shh* mRNA expression profiles in regenerating blastemas (4 cm animals, n = 9) and developing limb buds (n = 6). Signal intensity (mean ± SEM) as a function of absolute (left) or normalized (right) distance from the posterior boundary. Spatial normalization reveals near-identical profile shapes, demonstrating size-dependent scaling of SHH signalling.**g:** *Ptch1* gradient lengthscale as a function of blastema width across four animal size groups. Points show median ± IQR. A linear fit (blue dashed, animals ≤8 cm: λ = 105.8L + 0.4) and a square root fit (black dashed, all sizes: λ = 163.4√L − 54.6, R² = 0.963) are shown, indicating sublinear scaling of the SHH gradient with tissue size in larger animals. Fits are derived using the partially constrained background approach (see *Methods*). **h:** Left, simulated *Ptch1* gradient profiles with a single posterior source (blue) or an additional secondary anterior source (orange), showing a steeper endogenous gradient decay with ectopic SHH. Centre, log-transformed *Ptch1* mRNA signal intensity (mean ± SEM) in blastemas injected anteriorly with *Shh* + eGFP LNPs (blue, n = 8) or control eGFP LNPs (orange, n = 8), aligned at the position of minimum *Ptch1* amplitude (dashed line). Right, *Ptch1* gradient length scale is significantly shorter following *Shh* + eGFP LNP injection (Mann-Whitney p = 0.00466; n = 8 per group). Box plots show median and IQR.

To gain insight into possible mechanisms underlying such scaling behavior, we developed a minimal mathematical model for gradient formation and scaling of SHH signaling. In brief, we considered several defining features of the pathway, in particular that the receptor PTCH1 is both induced by SHH and promotes the degradation of SHH ^23–26^. A model based on these parameters alone does not display scaling and we therefore took the most minimal assumption of a biochemical expansion-repression mechanism (Fig. 2b) ^27,28^. In this model, an expander gene produces a protein that binds to the morphogen, reducing its degradation rate and/or enhancing its diffusion, thereby extending the typical morphogen length-scale. Expander expression is negatively regulated by morphogen signalling and the expander molecule is highly diffusible. These assumptions result in an expander protein distribution that is approximately uniform and increases with blastema size. We demonstrate analytically (*Supplementary theory*) and through numerical simulations that this mechanism can generate size-dependent scaling of SHH signaling in the SHH–PTCH1 system. Furthermore, consistent with our transcriptomic data, the model predicts robust scaling at small to intermediate blastema sizes, but a breakdown of scaling in larger tissues (Fig. 2c). This breakdown is attributed to the limited diffusion range of the expander protein (Fig. 2d), which limits its capacity to scale proportionally in larger systems.

This minimal model made a number of predictions on the relationship between *Shh* and *Ptch1* expression. However, testing these predictions required higher resolution quantification of *Shh* domain sizes and *Ptch1* expression gradients in different sized blastemas. For this, we performed HCR in situ Hybridization followed by lightsheet imaging of whole mount tissues. A first prediction of the model is an exponential decay profile of the *Ptch1* mRNA along the posterior-anterior axis, which could describe the data (Fig. 2e). A second prediction was that the Ptch1 decay length should increase with tissue size, but start plateauing above a critical size. We first evaluated whether we could detect scaling of the Ptch1 gradient when comparing developing limb buds to small blastemas from 4 cm animals. Although the *Shh* and *Ptch1* expression domains are larger in the blastemas, the amplitude of their expression resembles the one found in developing limb buds. After spatial normalization, the signaling gradients closely overlap (Fig. 2f), indicating proportional scaling of signaling between limb development and regeneration . We next asked whether gradient scaling breaks down at larger sizes. Comparing *Ptch1* gradient lengthscales across limb buds and blastemas from 4, 8, and 13 cm animals, we found a near-linear increase up to ∼2 mm tissue width (8 cm animals), beyond which the gradient length-scale began to saturate (Fig. 2g). Importantly, the same saturation was observed regardless of the fitting modality used to extract the Ptch1 lengthscale (*Methods*; Fig. S3a–d), and persisted when the lengthscale was plotted against animal size rather than blastema width (Fig. S3e). This was despite a continuous increase in size of the Shh domain (Fig. S3f), consistent with previous observations ^4^. As a third model prediction, the specific topology of the SHH-Ptch1 interaction network and degradation of PTCH1-SHH as a complex ^23,26^, predicts less intuitive spatio-temporal characteristics of the Ptch1 gradient. For instance, the model predicted a shift of the peak of the *Ptch1* gradient with respect to the anterior end of the *Shh* expression domain (Fig. ST2 in *Supplementary Theory*), which we could observe experimentally (Fig. 2e, Fig. S3a-c). We also found evidence that in subsequent timepoints in regeneration the *Ptch1* gradient amplitude increases and spatial range expands, while gradient length-scale decreases (Fig. S4a), reflecting progressive sharpening and broadening of the SHH signaling domain over time. Again, this was well recapitulated in the model (Fig. ST6 in *Supplementary Theory*). Altogether, this shows that our minimal model can recapitulate a number of complex features of Shh signalling gradients during regeneration.

Finally, we wished to test for the presence of an expander mechanism, feedbacking on SHH signalling range. After exploring computationally different possible perturbations, we found that a strong prediction is that ectopic activation of SHH signaling, distant from the endogenous source, should suppress local expander production and thereby reduce the length-scale of the endogenous *Ptch1* gradient. In the model, adding an anterior source of SHH robustly reduced posterior gradient length scale, by around 10-30% for realistic ectopic amplitudes (Fig. S4b). To test this prediction agnostic of expander identity, we used LNP-mediated mRNA delivery to express *Shh* in the anterior side of the blastema, and measured the endogenous *Ptch1* gradient after 48 hours. These experimental measurements showed a decrease in the lengthscale of the posterior-to-anterior *Ptch1* gradient (Fig. 2h), consistent with the model prediction of a reduction in production of an expander whose expression is negatively regulated by SHH signaling. Importantly, gradient amplitude was not significantly affected by this perturbation (Fig. 2h, Fig. S4b). Recent work has implicated Scube2 as a modulator of SHH spreading that is under negative regulation in the zebrafish neural tube ^29,30^. *Scube2* mRNA expression in the blastema was spatially uniform rather than complementary to the *Ptch1* gradient (Fig. S4c), suggesting the presence of an alternative or additional expander component unless *Scube2* is regulated post-transcriptionally .

These findings establish that SHH signaling exhibits feedback-mediated, size-dependent scaling during axolotl limb regeneration consistent with an expansion-repression mechanism, which breaks down in larger animals.

### Scaling of *Wnt9a-*modulated digit patterning

Next, we wanted to understand how scaling of SHH signalling and its failure in larger animals related to the subsequent stage of regeneration, digit patterning. The skeleton after lower arm regeneration was visualized via Alcian Blue/Aliziran Red staining (Fig. 3a, Fig. S5a). Quantification of digital elements revealed that digit number, as well as the number of metacarpals and phalanges was constant from small to mid-sized animals, but decreased for animals larger than ∼12cm including lack of an entire digit (Fig. 3b, Fig. S5b). While digit four was often missing, or appeared reduced and disconnected from the carpals in large animals, the number of phalanges decreased for all digits (Fig. S5b).

**Figure 3:**
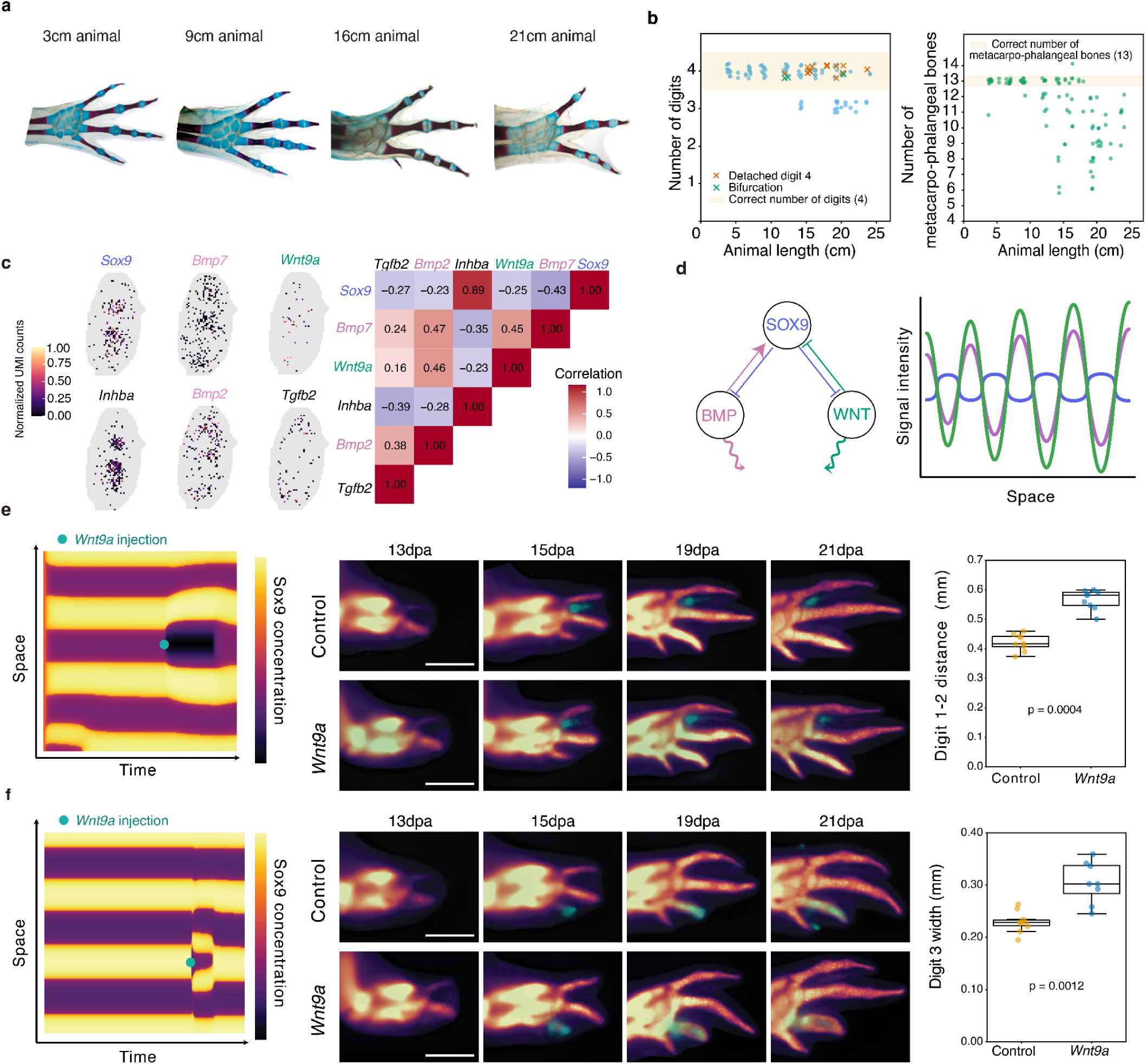
**a**. Regeneration fidelity decreases with animal size. Representative images of regenerated limbs from animals of increasing body length (3, 9, 16, and 21 cm), showing progressive loss of skeletal elements in larger animals. **b.** Quantification of regeneration fidelity across animal sizes. Left, number of digits plotted against animal length. Filled circles indicate standard regenerates with 3 or 4 digits; detached digit 4 (orange cross) and bifurcations (green cross) are labeled differently. Beige shading indicates the correct digit number (4). n= 117 limbs. Right, total count of metacarpo-phalangeal bones plotted against animal length; beige shading indicates the correct element count (13). Both metrics show reduced regeneration fidelity in animals larger than ∼10 cm. Each point represents an individual limb. **c.** Spatial gene expression and correlation analysis in the regenerating blastema. Left, spatial transcriptomics showing expression patterns of *Wnt9a*, *Inhba*, *Bmp7*, *Bmp2*, *Sox9*, and *Tgf-β2.* Color intensity (UMI counts) indicates expression level. Right, pairwise correlation matrix. *Inhba* shows positive correlation with *Sox9*, while *Wnt9a*, *Bmp7*, *Bmp2*, and *Tgf-β2* show negative correlations with *Sox9*. **d.** Reaction-diffusion model for digit patterning adapted from (Raspopovic 2014). Left, regulatory network: BMP activates SOX9, WNT inhibits SOX9, and SOX9 negatively feeds back on both pathways. Right, simulated spatial pattern showing periodic SOX9 peaks (blue) with WNT (green) and BMP (magenta) peaks out of phase, consistent with Turing instability and experimental spatial correlations. a.u., arbitrary units. **e**. *Wnt9a* overexpression increases interdigital spacing. Left, model prediction: transient WNT source in the interdigital region causes SOX9 domain repulsion. Middle, live imaging (13–21 dpa) of control (top) and *Wnt9a*-treated (bottom, LNP-delivered *Wnt9a* mRNA to interdigital space) limbs. Scale bars, 1 mm. Right, digit 1–2 spacing (measured at 19dpa) is significantly increased by *Wnt9a* (p = 0.0004). Box plots show median and IQR. *n* = 8-9 limbs per group. **f**. *Wnt9a* misexpression on a forming digit causes domain enlargement. Left, model prediction: transient WNT source placed on a SOX9 domain causes local splitting into two digit domains. Middle, live imaging (13–21 dpa) of *Wnt9a*-treated (bottom, LNP-delivered *Wnt9a* mRNA to digit 3) and control (top) limbs showing digit enlargement with central signal reduction. Scale bars, 1 mm. Right, digit 3 width (measured at 21dpa) is significantly increased by *Wnt9a* (p = 0.0012). Box plots show median and IQR. *n* = 8 limbs per group.

To understand this failure in digit formation, we sought to understand the molecular patterning mechanisms occurring in the blastema. Previous work in mouse limb development implicated a BMP-WNT-SOX9 Turing network regulating periodic digit specification ^31^. To identify candidate molecular players active in Axolotl digit patterning, we analyzed late timepoints of our spatial transcriptomics dataset focusing on spatial patterns that were in or out of phase with nascent digit condensations (*Sox9^+^* domains). *InhibinA* was found in phase with nascent digit regions while *Wnt9a, Bmp7, Bmp2* and *Tgfβ2* were enriched in the *Sox9^-^* interdigit regions (Fig 3c).

To evaluate the functional relevance of this pattern of gene expression on digit specification, we first adapted a reaction-diffusion model previously developed for mouse digit patterning ^31,32^, in which BMP activates SOX9, WNT inhibits SOX9, and SOX9 feeds back negatively on both pathways (Fig. 3d, left panel; *Supplementary Theory*). Across a broad range of parameters and in agreement with prior theoretical work ^33^, this yields a patterning instability with periodic digit formation, with the same spatial correlations that we observed experimentally (Fig. 3d, right panel; Fig. S5c,d). To challenge the model further, we tested whether it can predict specific perturbation responses. For instance, placing a transient ectopic source of WNT in an interdigit region in the model resulted in effective repulsion of the neighboring digit domains (Fig. 3e, left panel). On the other hand, placing this transient source in a digit region predicted a local splitting into two *Sox9*^+^ digit domains (Fig. 3f, left panel). To test these predictions, we used LNPs to deliver *Wnt9a* mRNA to the interdigital space between digits 1 and 2 or to the forming third digit. We performed this experiment in *Sox9-*eGFP transgenic reporter animals that allow live visualization of digit formation. With an interdigital source, *Sox9* domains consistently reconfigured, exhibiting effective repulsion from the *Wnt9a* source, resulting in a larger interdigital spacing (Fig. 3e, Fig. S5e). When the *Wnt9a* source was applied in the center of the forming third digit, we observed local attenuation and lateral expansion of *Sox9*, resulting in an enlarged digit three (Fig. 3f) with non-monotonous Sox9 expression profile (Fig. S5f). These dynamic responses closely match our numerical simulations, and constitute hallmarks of a dynamic patterning mechanism where diffusible species can act at a distance to drive digit spacing.

### Digit defects in large animals arise from growth failure

To relate the final digit outcomes seen in the Alcian Blue/Alizarin Red staining to the dynamics of digit formation, we performed live imaging of limb regeneration in *Sox9-*eGFP knock-in reporter animals of five different sizes, spanning from developing larvae to 21cm animals (Fig. 4a, Fig. S6a,b). Consistent with previous reports, we found that axolotl digits are specified progressively in the order 2-1-3-4 ^19^ and that, during digit specification, the tissue appears to expand. In any periodic patterning model on a growing domain, the number of digits is determined by the ratio between blastema width and the digit patterning length scale as well as on the amount of tissue growth. We first measured the relationship between patterning length scale at the 3 digit stage (the last common stage across animal sizes) and tissue width. We found that the distance between successive *Sox9*-positive digit condensations scaled with tissue width at all animal sizes (Fig. 4b). This is consistent with our spatial transcriptomics analysis of the blastema showing that WNT signaling and skeletal differentiation scale better than SHH signaling (Fig. 1f), and suggests that the digit patterning network scales even in large blastemas. This led us to probe whether differences in blastema growth in large animals could explain digit loss. Interestingly, we found that extensive tissue growth along the anterior-posterior axis (tissue width) occurs during the patterning process of small and medium-sized, but not large, animals (Fig. 4c, Fig. S6c). Since digits are added progressively, reduced growth would be expected to truncate the addition sequence, predominantly affecting the last-added fourth digit. Incorporating tissue growth in our simulations confirmed this intuition (Fig. 4d and *Supplementary Theory*).

**Figure 4:**
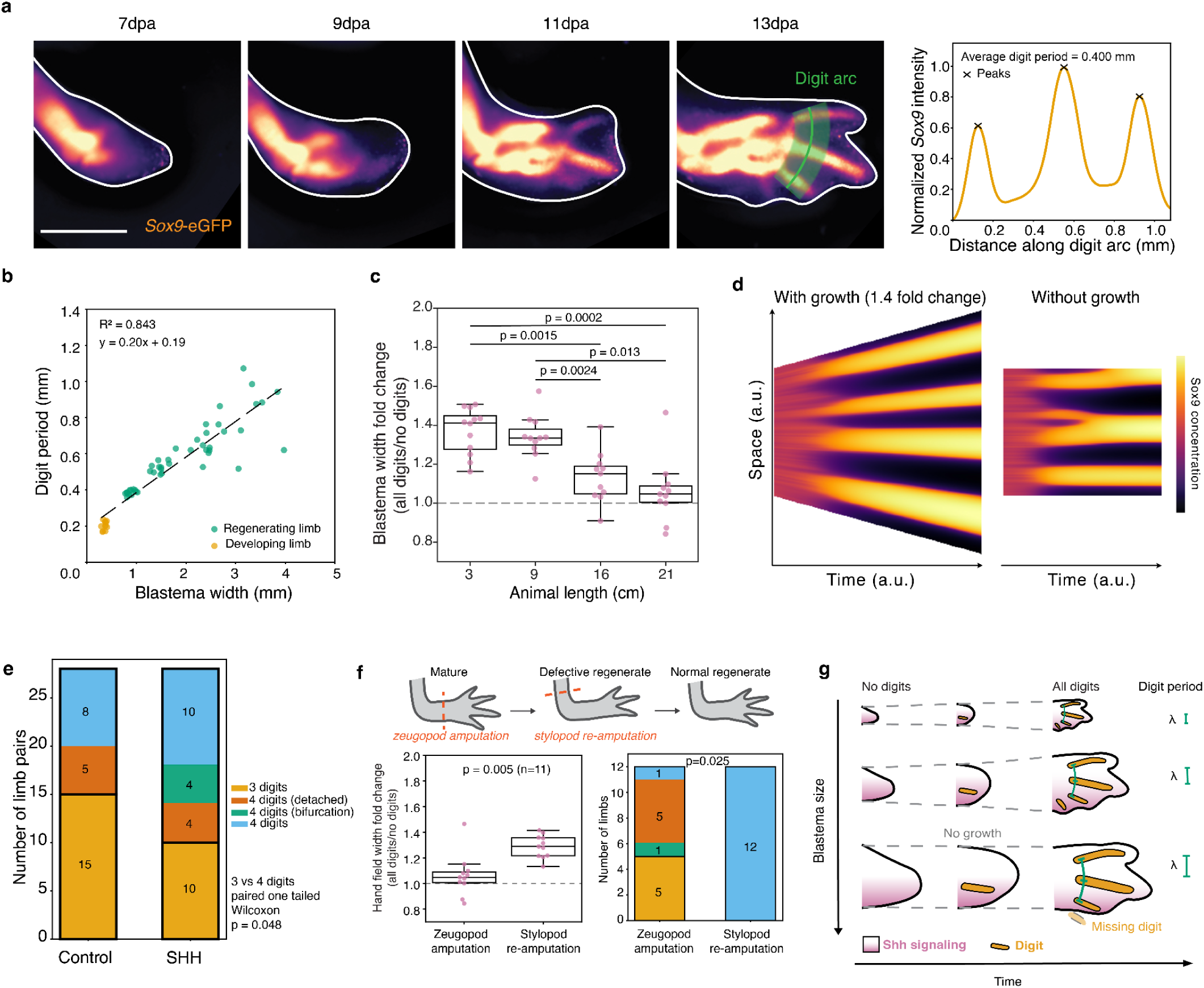
**a**. Time-course live widefield imaging of *Sox9*-eGFP expression in a regenerating limb (3 cm animal). High-intensity signal marks *Sox9*+ condensations. Right, quantification of *Sox9* intensity along the digit arc (green line, 13 dpa). Peaks correspond to digit positions; the average distance between consecutive peaks represents digit periodicity. Scale bar, 1 mm. dpa, days post amputation. **b**. Digit period scales with blastema width. Digit period plotted against blastema width (measured before digit emergence) in regenerating (orange) and developing (green) limbs. The dashed line indicates a linear fit, consistent with a size-scaled patterning mechanism. n= 57 limbs. **c.** Blastema width fold change during digit patterning across animal sizes. Each point represents the ratio of blastema width at the "all digits" stage (after digit series completion) to the "no digits" stage (before digit emergence) for an individual limb. Box plots show median, interquartile range, and whiskers extending to 1.5× IQR. Dashed line at fold change = 1 indicates no expansion. Horizontal bars indicate significant pairwise differences (Dunn’s post hoc test with Benjamini–Hochberg FDR correction following Kruskal–Wallis test). Blastema width increases significantly more during digit patterning in smaller animals (3 and 9 cm) than in larger animals (16 and 21 cm). n = 11-12 blastemas per group. **d.** Turing model simulation of digit patterning with and without domain growth. Left, simulation with 1.4-fold domain growth generates four digits. Right, simulation without growth leads to digit loss and pattern collapse, recapitulating the phenotype observed in large animals. **e.** Ectopic *Shh* delivery rescues digit numbers in large axolotls. Bar plot showing digit count in paired control and *Shh* mRNA-treated limbs (*n* = 28 limb pairs). Yellow, three digits (defective); blue, four digits (normal); green, digit 4 emerges as a bifurcation of digit 3; red,digit 4 is detached from carpals. SHH treatment significantly increases digit number compared to contralateral controls (one-tailed Wilcoxon signed-rank test, H₁: SHH > Control, *P* = 0.048). **f.** Stylopod re-amputation rescues digit numbers in large axolotls. Top, experimental schematic: animals with defective zeugopod regenerates were re-amputated at the stylopod level after regeneration completion. Bottom left, blastema width fold change is significantly greater following stylopod re-amputation than zeugopod amputation (Wilcoxon signed-rank test, p = 0.005), indicating restored blastema expansion. Colors indicate digit counts as in e . Box plots show median, interquartile range, and whiskers extending to 1.5× IQR. Dashed line at fold change = 1 indicates no expansion. *n* = 11 limbs. Bottom right, digit number is significantly increased following stylopod re-amputation compared to zeugopod amputation (Wilcoxon signed-rank test, p = 0.025, n = 12 limbs). **g.** Model for size-dependent digit loss. SHH signaling (pink gradient) scales with blastema size in small and intermediate tissues (top and middle rows), enabling proportional field growth and formation of all four digits. In large blastemas (bottom row), SHH signaling becomes insufficient, leading to saturated growth. The digit patterning mechanism scales correctly, but limited spatial expansion results in digit loss.

Given the established role of SHH in promoting tissue growth during limb development ^34–37^, we hypothesized that inadequate scaling of SHH-dependent growth in large blastemas may underlie the observed growth defects and resulting digit loss. Consistent with this interpretation, the digit defects predominantly affect the last appearing, posterior fourth digit and resemble those associated with insufficient Shh signaling in both mammalian and amphibian limb development ^38–40^. We first confirmed that enhancing SHH signalling expands blastema size. Both LNP-mediated *Shh* delivery (as in Fig. 2h) and SMO agonist (SAG) treatment increased blastema width in small blastemas (Fig. S7a). To directly test whether restoring SHH-mediated growth can rescue digit formation in large (18-24 cm animals), we used LNP-mediated delivery of *Shh* mRNA into regenerating blastemas. To avoid perturbing initial digit field establishment, *Shh* mRNA was delivered to the posterior side at late stages of regeneration (Fig. S7b), after the first digits had likely already formed- and when blastema growth was limited in large blastemas. Importantly, *Shh*-injected blastemas showed a higher proportion of regenerates with four digits (Fig. 4e). These results support the model that SHH-dependent growth is the limiting factor: locally augmenting SHH signaling can partially rescue digit formation in large blastemas.

An alternative explanation for missing digits is that the advanced age of larger axolotls may impair regenerative fidelity, independently of blastema size. In axolotls, the diameter of the upper arm is smaller than the lower arm, presenting a situation where we could interrogate a blastema with smaller field size on animals of the same age. We therefore amputated at the level of the upper arm the large animals that had previously regenerated defective digits. Strikingly, the resulting narrower upper-arm blastemas displayed rescued regenerative potential, with all samples faithfully regenerating four digits (Fig. 4f). Furthermore, as expected in our model, these upper-arm blastemas displayed similar growth as their blastema-width size matched lower-arm counterparts, confirming that this amputation location rescues both growth and digit formation (Fig. 4f, Fig. S7c). Accordingly, during the regeneration process upper arm blastemas produced smaller hands and closer-spaced digits than lower arm regenerates of the same animals. Altogether, these two rescue experiments argue in favor of a model where digit spacing scales with blastema size (Fig. S7d), but SHH-dependent growth fails to keep up in large lower arm blastemas, resulting in termination before regeneration of the last digit (Fig. 4g).

## Discussion

Here we took advantage of the remarkable regeneration capabilities of the axolotl to identify general principles, possible limitations, and restorative strategies for the regeneration of large adult tissues. While limb development in an individual species occurs at a fixed size, that spans only a 1-3 millimeters range across genetically different mouse ^41^, human ^42^, and whale ^43^ species, an individual axolotl is capable of regenerating limbs that are almost an order of magnitude larger than their developmental counterparts. As we show here, proportionate regeneration is not simply achieved by patterning at a fixed scale followed by expansion, but by scaling core signaling processes across tissues of very different sizes. At the same time, this scaling is not unlimited and patterning breaks down in the largest blastemas, revealing an upper physical limit to regenerative fidelity that results in digit loss. To understand these phenomena, we interrogated how different signaling and patterning events scale across a wide range of animal and tissue sizes during limb regeneration. Using spatial transcriptomics, quantitative imaging and functional assays we found that signal-dependent growth and patterning scale in an increasingly different manner with blastema size. This system yields accurate patterning across most animal sizes but breaks down in the largest animals, where discordant scaling of growth and patterning results in imperfect regeneration.

We observed significant scaling of SHH signaling, as evidenced by *Ptch1* gradient lengthscales expanding up to fourfold, a far larger extent than what has been observed in developmental systems. However, this scaling saturates in the largest animals, where both the SHH signaling domain and tissue growth become sublinear with blastema size. Using biophysical modelling we show that this behaviour can be predicted by a size dependent feedback mechanism. *Ptch1* gradient scaling had not been observed by Furukawa et al ^4^, likely due to a combination of confounding factors, including their use of primarily larger sized animals where subscaling is observed, the use of colorimetric methods that are inherently non-linear and may lack sensitivity. Lyubaykina et al ^5^, through theoretical approaches, predicted body-size-dependent static scaling of two opposing morphogen gradients as a means for achieving an appropriate final size. Here, we observe dynamic scaling of SHH-signaling, that is responsive to local perturbations and followed by cell patterning and differentiation. Blastema size continuously increases during the course of regeneration where cell proliferation is controlled in several phases of signaling. Initially, induction of SHH and FGF8 expression occurs in response to injury but then the maintenance of their expression becomes mutually dependent. In the midst of this second growth phase, cell specification and patterning of cartilage is established, under the influence of a periodic patterning network. We find that digit periodicity scales linearly, while tissue growth and SHH signaling do not. This means that digit spacing becomes mismatched compared to the evolving size of the blastema, precluding or altering the emergence of the final, posterior digit. This phenotype, analogous to those associated with insufficient Shh signaling in both mammalian and amphibian limb development^34–37^, emerges in the absence of genetic or pharmacological perturbations, exposing a limitation of the endogenous patterning systems. While our analysis focused on the spatial extent of SHH signaling, reduced signaling range could lead to earlier termination of SHH expression, as the mutual maintenance of SHH and FGF8 expression depends on sufficient signaling overlap. This possibility is difficult to examine directly, as larger animals require longer times to complete regeneration. However, the defects we observe are not restricted to posterior digits but extend to phalangeal loss in anterior digits, suggesting that signaling range itself plays an important role. Importantly, these distinct aspects of signaling converge on proliferative expansion of the patterning field, which has been proven sufficient to restore the digit number even upon defective SHH signaling ^44^. After the digit patterning phase, complete restoration of limb proportionality, especially along the proximal distal axis, requires a final phase of post-patterning growth ^19^.

Interestingly, two extreme periodic patterning scenarios which have been discussed in the past are either a classical Turing patterning model, where wavelength is independent of domain size, or perfectly scaling patterns, where the wavelength grows linearly with domain size ^28,45,46^. In the context of digit patterning, the former predicts an increasing number of digits with domain size ^47^, while the latter predicts a constant digit number. Our observations reveal a third and qualitatively different outcome, in which digit number decreases with animal size. This emerges because tissue size influences digit patterning at multiple levels: digit periodicity scales with blastema size at the onset of patterning, such that blastemas of different sizes begin with appropriately scaled wavelengths, while continued growth during the patterning phase allows the sequential addition of further digits. Final digit number therefore depends on the dynamic interplay between this scaled wavelength and ongoing domain expansion, and breaks down when SHH-dependent growth saturates in the largest blastemas. Importantly, this framework does not exclude continued scaling of the factors that set digit periodicity within an individual growing blastema. Even in that case, posterior-biased growth together with pattern locking of already-specified digits could account for sequential digit addition, as previously placed digits would remain fixed while newly available tissue accommodates additional ones ^48–50^. Importantly, this mechanism of concordant or discordant scaling in patterning wavelength versus growth of domain size is not specific to Turing-like reaction diffusion mechanism, but should hold generically in other contexts for any mechanisms yielding a period patterning instability, including purely mechanical patterning mediated by cellular contraction ^51,52^ or more complex models involving feedbacks between mechanics and signalling ^53,54^ .

Overall, our work provides technical advances connecting spatial-omics with concepts from morphogen scaling ^7,55–57^ and pattern formation ^58,59^. Measuring properties such as domain scaling, signalling gradients and spatial correlations, we use the spatial information to infer underlying biophysical mechanisms of morphogen reaction-diffusion dynamics.We furthermore provide conceptual insight into future regenerative engineering of adult tissues. Importantly, size dependent failures in spatial patterning have recently been observed in mouse gastruloids ^60^ and in *ex vivo* presomitic mesoderm cultures ^61^, outlining the broader relevance of our findings. As signaling pathways appear to have independent scaling mechanisms, achieving careful coordination of scaling among several pathways will be required to avoid serious defects. Synthetic biology and engineering approaches will be interesting ways to face this challenge.

## Supporting information

Supplementary Theory

## Acknowledgements

We sincerely thank the STOmics Grant Program for granting us access to the Stereo-seq technology and the STOmics EU team for their continued support of this project. We thank all members of the Hannezo and Tanaka labs for discussions, in particular Katharina Lust and Diego Rodriguez Terrones. We acknowledge Lorenzo Magrini for advice on data fitting. We thank the IMBA-IMP Bio-optics facility (especially Alberto Moreno Cencerrado, Pawel Pasierbek and Gabriele Bradamante), IT, Ethics and Amphibian caretaking service staff. We thank Benjamin Friedrich and Natalia Lyubaykina for discussions and input early in the project.

## Use of artificial intelligence

AI tools were used for coding assistance and text editing. All outputs were reviewed and verified by the authors.

## Author contributions

The project was conceptualised by PT, EMT, FO and EH. Experimental work was performed by PT together with MNA (digit patterning perturbations, skeletal stainings), DK (spatial sequencing), JL (Scube2 analysis) and YTS (skeletal stainings). FO analyzed spatial sequencing and developed theoretical models and numerical simulations with input from EH, PT and EMT. Image data analysis was performed by PT with input from EMT, FO and EH. MNA and JL quantified skeletal preparations. Data was interpreted by PT, FO, MNA, EMT and EH. AF provided protocols and support for LNP preparation. PM generated the transgenic *Sox9-*eGFP line. The manuscript was written by PT, EMT, FO and EH. EMT and EH provided supervision and secured funding. All authors approved this manuscript.

## Methods

### Animal housing and surgical procedures

Homozygous white (*d/d*) and transgenic axolotls were bred and maintained at the IMP/IMBA animal facilities. All animals were housed individually. Experimental procedures were conducted in accordance with local animal ethics committee guidelines and approved by the Magistrate of Vienna (animal licenses: GZ: 51072/2019/16, GZ: MA58-1432587-2022-12, GZ: MA58-1516101-2023-21, 2024-0.438.721 and 2024-0.438.718). Limb amputations were performed bilaterally on the forelimbs of all animals. Unless otherwise specified, amputations were made through the mid-zeugopod (lower arm). Stylopod re-amputations (Fig. 4f) were performed through the mid-stylopod (upper arm).

### Transgenic lines

All lines used in this study are in the *d/d* white mutant background, the commonly used laboratory strain. The *Hoxa13* reporter line *tm(Hoxa13t/+:Hoxa13-T2A-mCherry)^Etnka^* line has been previously described ^1^. The *tm(Sox9t/+:Sox9-eGFPnls-T2A-ERT2-Cre-ERT2)^Etnka^*line was generated for this study using the same CRISPR/Cas9 strategy and gRNA (GGACTGCTGGCGAATGCACC) as for the previously published *(Sox9:Sox9-T2a-mCherry)^Etnka^*line ^2^, with the following targeting cassette: eGFPnls-T2A-ERT2-Cre-ERT2 (Addgene plasmid #256056), resulting in eGFPnls fused to the C-terminus of *Sox9*, followed by a T2A self-cleaving peptide and ERT2-Cre-ERT2. Double transgenic animals were generated by intercrossing the *tm(Hoxa13t/+:Hoxa13-T2A-mCherry)^Etnka^*and *tm(Sox9t/+:Sox9-eGFPnls-T2A-ERT2-Cre-ERT2)^Etnka^* lines.

### Pain management

Prior to surgery, live imaging, or tissue harvesting, axolotls were anaesthetised by immersion in 0.01-0.03% benzocaine (Sigma) solution until unresponsive to physical stimuli. Following imaging, animals were returned to their housing tanks to recover. Following surgery, animals were maintained in butorphanol solution (0.5 mg/L) for 72 hours post-operatively, with a full solution exchange at 24 hours. Animals were euthanised by immersion in a lethal concentration of benzocaine solution at the experimental endpoint.

### Spatial transcriptomics

Spatial transcriptomics was performed on regenerating forelimbs from axolotls of three body sizes at three matched blastema stages (mid-stage, conical, palette). Tissue sectioning blocks were assembled containing one blastema from 20 cm animals, two blastemas from 12 cm animals, and 2–3 blastemas from 4 cm animals. Blocks were prepared to represent two blocks per stage (6 blocks total). From each block, one tissue section was collected onto a Stereo-seq chip, except for two blocks from which two sections were collected, resulting in 8 chips in total. Sectioning blocks were prepared as follows: harvested samples were washed in 0.7× PBS, transferred directly to optimal cutting temperature (OCT) compound, and flash-frozen in isopentane chilled with liquid nitrogen. Samples were stored at −70°C until sectioning. All subsequent steps, including cryosectioning, Stereo-seq library preparation, sequencing, and raw data processing, were performed at the BGI facility (Riga, Latvia) in collaboration with BGI, supported by a STOmics Grant awarded to E. Tanaka. The Stereo-seq protocol was performed as previously described ^3^, with the following modifications: tissue sections were cut at 20 µm thickness and permeabilised for 6 minutes using 0.5× permeabilisation reagent.

### HCR in situ hybridization and tissue clearing

For HCR in situ hybridization, developing limb buds and regenerating blastemas were collected from animals of defined size groups. For the limb bud-to-blastema comparison (Fig. 2e,f), limb buds were obtained from animals of ∼1.7 cm nose-to-tail length, and blastemas from ∼4 cm animals. For the comparison across multiple sizes, blastemas were collected from ∼1.7, ∼3.0, ∼4.0, ∼8.0, and ∼13.0 cm animals at stage-matched timepoints corresponding to the mid-bud blastema. Developing limb buds were collected at stage 45-47. Samples were processed as previously described ^4^. Harvested limbs were fixed in 4% paraformaldehyde (PFA) at 4°C overnight (14–17 hr), then progressively dehydrated through a methanol/PBS series (25:75, 50:50, 75:25, and 100:0 methanol:PBS), each step on ice for a minimum of 30 min. Samples were stored in 100% methanol at −20°C until use. Prior to staining, samples were rehydrated through the reverse methanol/PBS series (25:75, 50:50, 75:25, and 0:100 methanol:PBS). Probes were designed against *Shh*, *Fgf8*, *Ptch1*, and *Scube2* as previously described . Probes for *Shh, Fgf8* and *Ptch1* were purchased from Molecular Instruments; probes for *Scube2* were ordered as oligopools (IDT Opools). HCR in situ hybridization and tissue clearing were performed as previously described^4,5^. Briefly, stained whole-mount samples were cleared in Ce3D solution and imaged on a Z1 lightsheet microscope.

### LNP preparation and injection

Lipid nanoparticles (LNPs) were prepared as previously described^6^. Briefly, SM102-based LNPs were formulated at an N/P ratio of 6.0, washed with PBS using Amicon Ultra 10 kDa centrifugal filters (Merck), and diluted to a final mRNA concentration of 1.5 × 10⁻⁷ mol/L. Encapsulation efficiency (EE%) was determined using the Quant-iT™ RiboGreen RNA Assay Kit (Invitrogen); only LNPs with EE% ≥ 90% were used for injections. LNPs were delivered using a Nanoject II injector (Drummond Scientific Company).

Four LNP preparations were used in this study. For *Shh* misexpression, LNPs encapsulated *Shh* mRNA and *GFP* mRNA at a 1:1 molar ratio; the corresponding control LNPs contained equimolar amounts of *GFP* mRNA alone. To generate an ectopic anterior *Shh* source (Fig. 2h), 4–9 nl of LNP preparation was injected per blastema; for rescue experiments in larger animals, this volume was increased to 23 nl. For *Wnt9a* misexpression, LNPs encapsulated *Wnt9a* mRNA, BlueBonnet mRNA, and UTP-PEG5-AZDye647-labelled mRNA at a 4:4:1 molar ratio; the corresponding control LNPs contained BlueBonnet and UTP-PEG5-AZDye647-labelled mRNA at a 9:1 molar ratio. For all *Wnt9a* experiments, 4 nl of LNP preparation was injected per animal. The 647-conjugated mRNA encodes BFP but is not expected to produce functional protein, serving solely as a fluorescent tracer to confirm LNP delivery. UTP-PEG5-AZDye647-labelled mRNA was generated by *in vitro* transcription using the HighYield T7 AZDye647 RNA Labeling Kit (UTP-based; Jena Bioscience, cat. no. RNT-101-AZ647).

### Wnt9a misexpression

*Wnt9a* misexpression was performed in 5–6 cm long double transgenic axolotls (*Hoxa13-T2A-mCherry*; *Sox9-eGFPnls*). Ten animals were injected with *Wnt9a* LNPs and ten animals received control LNPs. One *Wnt9a*-allocated animal was excluded from the study due to illness unrelated to the experimental procedure. Injections were performed at 13 days post-amputation, when digit 1 and digit 2 were first visibly separate. In each animal, one limb was injected in the interdigital space between digit 1 and digit 2, and the contralateral limb was injected in the region of the forming digit 3 with the same LNP preparation. Successful LNP delivery was confirmed by live imaging of the AZDye647 fluorescent tracer. Injected limbs were imaged every two days from 13 to 23 dpa, and again at 29 dpa.

### Anterior Shh misexpression

*Shh* anterior misexpression was performed in 4-5cm d/d animals. Twelve animals were injected with the *Shh* LNPs and 11 animals received control LNPs. Injections were performed in the anterior side of the blastemas 7 days post amputation (conical stage blastemas). Limbs were live imaged and then harvested for HCR in situ hybridization 2 days post injection, from 6 *Shh* injected animals and 6 GFP injected animals. The remaining animals were allowed to regenerate.

### Posterior Shh misexpression

Posterior *Shh* misexpression was performed in 18–24 cm animals. Animals were randomly assigned to one of three experimental groups: (1) *Shh*+*eGFP* LNPs versus *eGFP* LNPs (contralateral limbs of the same animal), (2) *Shh*+*eGFP* LNPs versus uninjected, and (3) *eGFP* LNPs versus uninjected. Injections were performed at 21 dpa, corresponding to early digit formation. Limbs were imaged every two days from 21 to 37 dpa. Samples were harvested for Alcian blue and Alizarin red staining following regeneration completion (78 dpa).

### Alcian blue and Alizarin red staining

Skeletal staining was performed on fixed limbs following a previously described protocol^7^. Limbs were harvested from both transgenic *Sox9-*eGFP and d/d animals (Supp. table). Rare instances of complete lack of regeneration were excluded from the dataset. Cartilage and bone were visualised using Alcian blue and Alizarin red staining, respectively. Stained limbs were imaged in 100% ethanol using an Axiozoom V16 Stereomicroscope (Zeiss) with an AxioCam 712 color camera (Zeiss) or an Olympus SZX10 stereomicroscope with an AxioCam ERc 5s colour camera (Zeiss). Alcian blue (cat. no. A3157), Alizarin red (cat. no. A5533), and Trypsin (cat. no. 85450C) were purchased from Merck.

### Skeletal elements quantification

Phalangeal, metacarpal, and carpal elements were counted manually by the same experimenter for all specimens. Digit number was scored as 4 in three cases: when four metacarpals were present; when three metacarpals were present alongside at least one elongated cartilage element in the putative position of digit 4, interpreted as a detached or incompletely separated metacarpal; or when a fourth digit tip arose as a bifurcation from digit 3 (Supplementary Fig. 5a). In all other cases with three metacarpals and no additional elongated element, digit number was scored as 3. Small, non-elongated spot-like condensations were not counted.

### Live imaging of Sox9 expression in different size animals

Live imaging of *Sox9-eGFPnls* was performed in axolotl larvae during forelimb development and at 4 animal sizes during forelimb regeneration following mid-zeugopod amputation. All animals were from the same breeding; 6 animals were imaged at each size. Body lengths at the time of amputation were 3.1 ± 0.2, 8.9 ± 0.3, 16.4 ± 1.1, and 21.2 ± 1.9 cm (mean ± SD, measured nose to tail from widefield images). Animals were imaged every two days from prior to digit emergence until digit series completion: 3 cm animals from 7 to 15 dpa, 9 cm animals from 10 to 22 dpa, 16 cm animals from 11 to 22 dpa, and 21 cm animals from 17 to 35 dpa. Imaging was performed at 40× (3 and 9 cm animals), 20× (16 cm), and 16× (21 cm) magnification. All limbs were harvested for skeletal staining after regenerative patterning completion.

### Stylopod re-amputation of large animals

The 21 cm animals described above were re-amputated at the mid-stylopod level following regenerative patterning completion (69 dpa). At re-amputation, animals measured 21.2 ± 1.8 cm (mean ± SD). Regenerating limbs were imaged at 16× magnification every second day from 17 to 33 dpa, and at 37 dpa. Limbs were harvested for skeletal staining after regenerative patterning completion.

### Microscopy

#### Lightsheet imaging

Cleared whole-mount samples were imaged on a Zeiss Lightsheet Z.1 microscope using a ×20/1.0 Clr Plan-Neofluar Corr DIC detection objective (nd = 1.53, WD = 6.4 mm), with the correction collar adjusted to match the refractive index of the clearing medium (1.501). Illumination was provided by ×10/0.2 objectives. HCR probes were excited at 647 and 561 nm, autofluorescence was visualized using 488 nm excitation, and DAPI was excited at 405 nm.

#### Widefield imaging

Widefield images were acquired on a Zeiss AxioZoom V16 stereomicroscope equipped with a PlanNeoFluar Z 1.0×/0.25 objective (56 mm working distance) and a Hamamatsu ORCA-Flash 4.0 sCMOS camera. Magnification and exposure time were adjusted per experiment to accommodate specimen size and optimize signal quality, then held constant throughout each time-course.

#### Image analysis

All image analysis was performed using custom Python workflows. All code will be made available upon acceptance.

#### Analysis of whole mount lightsheet HCR images

Three-dimensional light-sheet microscopy images containing *Shh*, *Ptch1*, and autofluorescence channels were processed using a custom Python pipeline. Images were first downscaled to an isotropic voxel size of 3 µm. Inter-channel registration was applied where needed. Three pre-trained GPU-accelerated pixel classifiers (APOC/OpenCL) were used to segment the tissue boundary, the *Shh*-expressing domain, and blood vessels from the autofluorescence channel; where necessary, *Shh* domain segmentations were manually corrected. The epidermis was removed by morphological erosion of the tissue mask by a user-defined thickness, yielding a mesenchyme-only mask. Camera background was estimated as the median intensity of voxels outside the tissue and subtracted from all channels. Blood vessel voxels were excluded, producing a final composite mask (mesenchyme minus blood vessels) applied to all channels. Images were rotated in 3D (assessed interactively using pyclesperanto on GPU) to align the proximo-distal tissue axis with the image y-axis and the posterior-anterior axis with the image x-axis. Rotated volumes were cropped around the centre of mass of the *Shh* domain. One-dimensional anterior-posterior intensity profiles were obtained by averaging intensities along the z and y axes, retaining only positions where at least 25% of voxels contained valid (non-masked) data. Two-dimensional average-intensity projection maps were generated from the rotated masked volumes. All per-image parameters were stored in JSON files for reproducibility. For *Ptch1*/*Scube2* co-expression analysis, images were averaged in z across the entire blastema mesenchyme.

#### Measurements of digit 1 to digit 2 distance (*Fig. 3e*)

For each blastema, anchor points were placed along the base of the digits and a spline curve was fitted through these points. Signal intensity was sampled along a band of fixed width extending distally from the spline, perpendicular to the digit base. The intensities across the band width were averaged and projected onto the central spline to generate a single mean intensity profile per blastema. Profiles, coordinates, and metadata were saved alongside quality-control overlay images. The central mean intensity profile of each sample was smoothed using a Savitzky-Golay filter, and peaks were detected with predefined parameters. The distance between the first two peaks was used as a proxy for digit 1 to digit 2 spacing.

#### Measurements of digit 3 width (*Fig. 3f*)

For each blastema, two user-defined points were placed to define a line perpendicular to digit 3 through the injected area. Signal intensity was sampled along a band of fixed width centred on this line, and intensities across the band were averaged to produce a central mean intensity profile. The profile was smoothed using a Savitzky-Golay filter, and digit 3 width was measured as the distance between the left and right crossings of a fixed intensity threshold around the detected peak. Profiles, coordinates, and metadata were saved alongside quality-control overlay images.

#### Measurements of digit period (*Fig. 4b*)

For each blastema, measurements were taken at a user-defined timepoint at which digit 1, 2, and 3 appeared clearly separated. Anchor points were placed along the base of the digits and a spline curve was fitted through these points. Signal intensity was sampled along a band extending distally to the length of digit 3, perpendicular to the digit rays, and intensities across the band width were averaged to produce a central mean intensity profile per blastema. The profile was smoothed using a Savitzky–Golay filter and peaks were detected with predefined parameters. Digit period was calculated as the mean of the digit 1-digit 2 and digit 2-digit 3 inter-peak distances. Profiles, coordinates, and metadata were saved alongside quality-control overlay images. For scaling analyses, digit period was plotted against blastema width measured at the pre-patterning ("no-digits") stage, prior to digit specification, rather than at the time of period measurement.

#### Blastema width measurements

Blastema width was measured from limb images by manually drawing a straight line perpendicular to the proximodistal axis of the blastema. For early stages (prior to digit emergence), the line was placed at the base of the blastema. For later stages, it was placed at the base of the hand field, defined as the proximal boundary of Sox9 expression at the level of the forming carpals, just distal to the forming radius and ulna. For each image, a tissue mask was generated interactively using SAM-assisted labelling in Napari, and the length of the line contained within the mask was taken as the blastema width. For the *Sox9*-eGFP time-course experiment (Fig. 4c), blastema width was measured at two defined stages, identified by manual inspection of the time-course images: prior to digit emergence, and when the complete digit series was first present. For animals that did not regenerate four digits, the final-stage measurement was taken 4 days after the three-separated-digits timepoint. A detached Sox9-positive element was considered equivalent to a fourth digit for the purpose of stage assignment. For all other datasets, blastema width was measured at a single stage corresponding to the relevant experimental timepoint.

### Exponential fitting of intensity profiles

For each one-dimensional intensity profile, the fitting window was determined automatically on the smoothed profile derivative: the fit start was placed at the first sustained negative derivative reaching a fixed fraction of the steepest decline. The fit end was set where the derivative became persistently positive over a specified distance threshold, to avoid the far-tail rise or noise. Three exponential models were then fitted to the raw (unsmoothed) profile within this window by non-linear least squares. The free-background fit, *y* = *a* e^(−*x*/λ) + *c*, treated all three parameters as free. The fixed-background fit used the same form but fixed *c* to the 3rd percentile of the last 50% of the profile. The partially constrained background fit restricted *c* to within ±5% of the profile’s dynamic range around the minimum. For the anterior *Shh* misexpression experiment (Fig. 2h), additional constraints were applied to ensure comparable fitting between conditions. The fit window of eGFP control profiles was restricted to match the average fit length of *Shh*-injected profiles, since the latter were truncated by the secondary anterior source. In addition, the background was fixed to a common value across all fits, estimated as the global background intensity from control samples (indicated by the dashed horizontal line in Fig. S4b).

### Shh domain size measurements

We defined the SHH domain width from the 1D SHH intensity profile by smoothing, subtracting a background estimated from the low-percentile tail, and measuring the distance between the left and right crossings at a fixed fraction (10%) of the peak intensity . The SHH volume was extracted directly from 3D segmentation (see image analysis section).

### Statistical analysis

The following details statistics for key experiments shown in the main figures. Statistical details for all other experiments are reported in the corresponding figure legends.

#### Wnt9a interdigital overexpression increases interdigital spacing (*Fig. 3e*)

Digit 1-digit 2 distances were compared between Wnt9a-injected and control animals using a two-sided Mann-Whitney U test (n = 8-9 per group). Control: mean = 421.8 µm, SD = 28.5 µm; Wnt9a-injected: mean = 566.8 µm, SD = 33.0 µm; U = 81.0, p = 0.0004.

#### Wnt9a misexpression in the forming digit 3 increases its width (*Fig. 3f*)

One *Wnt9a*-injected animal was excluded from analysis due to absence of LNP delivery within the digit. Digit 3 width was compared between Wnt9a-injected and control animals using a two-sided Mann-Whitney U test. Control: n = 10, mean = 229.3 µm, SD = 19.5 µm; Wnt9a-injected: n = 8, mean = 304.7 µm, SD = 39.9 µm; U = 77.0, p = 0.0012.

#### Blastema width fold change during digit patterning across animal sizes (*Fig. 4c*)

For each blastema, blastema width growth was expressed as the fold change in width from the no-digits stage to the final stage (all-digits / no-digits). The no-digits value was taken from the latest available pre-digit timepoint, and the all-digits value from the earliest four-digit timepoint; for animals that did not regenerate four digits, the final-stage measurement was taken 4 days after the three-separated-digits timepoint. Fold changes were compared across the four size groups (3, 9, 16, and 21 cm) using a Kruskal–Wallis test (H = 23.755, p = 2.8 × 10⁻⁵). Pairwise comparisons were performed using Dunn’s post hoc test with Benjamini-Hochberg false discovery rate correction: 16 cm vs. 3 cm, p = 0.0024; 16 cm vs. 9 cm, p = 0.013; 21 cm vs. 3 cm, p = 0.00025; 21 cm vs. 9 cm, p = 0.0015; 16 cm vs. 21 cm, p = 0.48; 3 cm vs. 9 cm, p = 0.58.

#### Shh rescue experiment (*Fig. 4e*)

Statistical analyses were performed on paired contralateral limbs. Animals whose Shh-injected limb showed no GFP expression at 4 days post-injection were excluded from the analysis. Non-injected and GFP-injected controls were first compared using a paired Wilcoxon signed-rank test; as no significant difference was detected (p > 0.05), both groups were pooled into a single control group for subsequent analyses. The pooled control group was then compared against Shh-injected limbs using a one-tailed paired Wilcoxon signed-rank test (H₁: Shh > Control; W = 35.0, p = 0.048). Of 28 paired animals, 9 were informative (discordant pairs); among these, 7 showed digit gain in the Shh-injected limb and 2 showed digit loss. Among the 15 animals with 3 digits in the control limb, 7 (47%) regenerated 4 digits in the Shh-injected contralateral limb. Of these 7 rescued limbs, 4 were complete and 3 were partial rescues presenting a detached or bifurcated fourth digit.

### Analysis of sequencing data

In this method section we give a summary on the statistical methods used for the analysis of spatial transcriptomic data in support of the main text. The code for reproducing all the sequencing panels is given as a separate tile. All the statistical analysis was performed in R with Seurat 5.0.0.

### Multiple-batch integration and clustering

In this section we give details on the statistical methods used for the inference of batch-corrected data where all the different blastemas in all the batches are analysed together in a single count matrix. Elements in the count matrix are given at a bin size of 50 μm resolution. This means that each bin does not correspond to a single cell, but transcripts from multiple cells might contribute in the same bin. After all the data sets are loaded we use the function SplitObject on the merged list of rds files, we then apply SCTranforms for the individual batches and the function SelectIntegrationFeatures with the first 1000 HvGs. We then use the functions PrepSCTIntegration and FindIntegrationAnchors with the SCT method to integrate all the batches. We check after UMAP dimensional analysis that the batches are well integrated as they should not be clustered in the lower dimensional space. On the integrated data we then performed the PCA, clustering and visualised the results on UMAP coordinates (Supp. Fig. 1b). In Supp. Fig. 1a we provide the total UMI counts distribution for each individual batch and blastema. We used the FindCluster function in Seurat to identify the epithelial population, Supp. Fig. 1d’ which is removed from further analysis.

### Scaling of gene expression

To quantify spatial patterning of gene expression, we selected the top 1000 highly variable genes (HVGs) from the integrated assay. For each gene, and separately within each blastema, we computed a Ripley-based clustering index. Specifically, for each gene, expression values across spatial locations were thresholded to identify highly expressing spots (top quantile of expression within each blastema). These spots were then treated as a spatial point pattern, and Ripley’s L-function was computed using translation correction. The patterning index was defined as the average deviation L(r) - r. This index quantifies the degree of spatial clustering: positive values indicate spatial aggregation (i.e. coherent expression domains), while values close to zero correspond to spatially uniform or random expression. Genes with positive Ripley index values were interpreted as exhibiting spatially structured expression patterns. To quantify domain size independently of the clustering metric, expression domains were also defined using an adaptive threshold (Otsu method) within each blastema; this yielded qualitatively consistent scaling relationships.

We then order genes based on their average patterning index across blastemas and in Fig 1d. And Fig. S1c we show the estimated domain as a function of blastema size for a different number of chosen genes. GO analysis in Fig 1d (right). was performed in R with the biomart package on the set of genes chosen for the scaling Fig 1d (left) after normalization. In order to measure scaling of identified clusters, we computed the number of bins for each cluster in each blastema. We then select only the clusters that are present in at least one blastema for each macro-category (small, medium and large). We then average these clusters for each blastema size and the resulting scaling is shown in Fig. 1e where the p-value and exponents are computed with a linear regression on the log-log transformed data. We finally performed GO analysis for the group of genes which are downregulated from small to large sized blastemas (Fig. 1f) and Fig. S1e using the FindMarkers() function in Rstudio with min_pct = 0.10, logfc.threshold = 0 and test.use = "wilcox". In Fig. 1f we subset 100 genes for visualization only, but the function was run on the top 1000 genes found from FindMarkers().

### Scaling of signaling pathways

In order to demonstrate how different pathways involved in limb-regeneration scale across blastema sizes, we computed the scaling of domains for a subset of genes belonging to each pathway. As for many genes we had few sparse counts, in order to avoid overestimating the domain size we defined the domain as the number of bins with a positive number of UMI counts for each gene. We then averaged across macro-category (small, medium and large) and the results are shown in Fig. S1f for each individual gene. The slope (a->b) is computed simply as the linear regression coefficient of the line passing through the two macro-categories a and b.

### Correlation matrix between genes

In order to compute the correlation matrix in Fig. 3c and Fig. S5c we first identified the batch which had SOX9 expression consistent with digit initiation. We found that the most reliable one in terms of average expression was the batch C1.

We then develop a pipeline to define 1D stripes and computed 1D gene expression profiles. For each blastema, we define stripes which pass through the weighted center of mass for SHH and FGF8 domain. We then projected individual bins inside the stripe on the midline of the stripe and averaged for equally spaced bins.

We then compute the Pearson correlation between pairs of 1D gene expression profiles and the results are shown in Fig. 3c for the selected genes and Fig. S5c for the first HVGs of the selected blastema (medium sized of the batch C1).

---

**Supplementary Figure 1:**
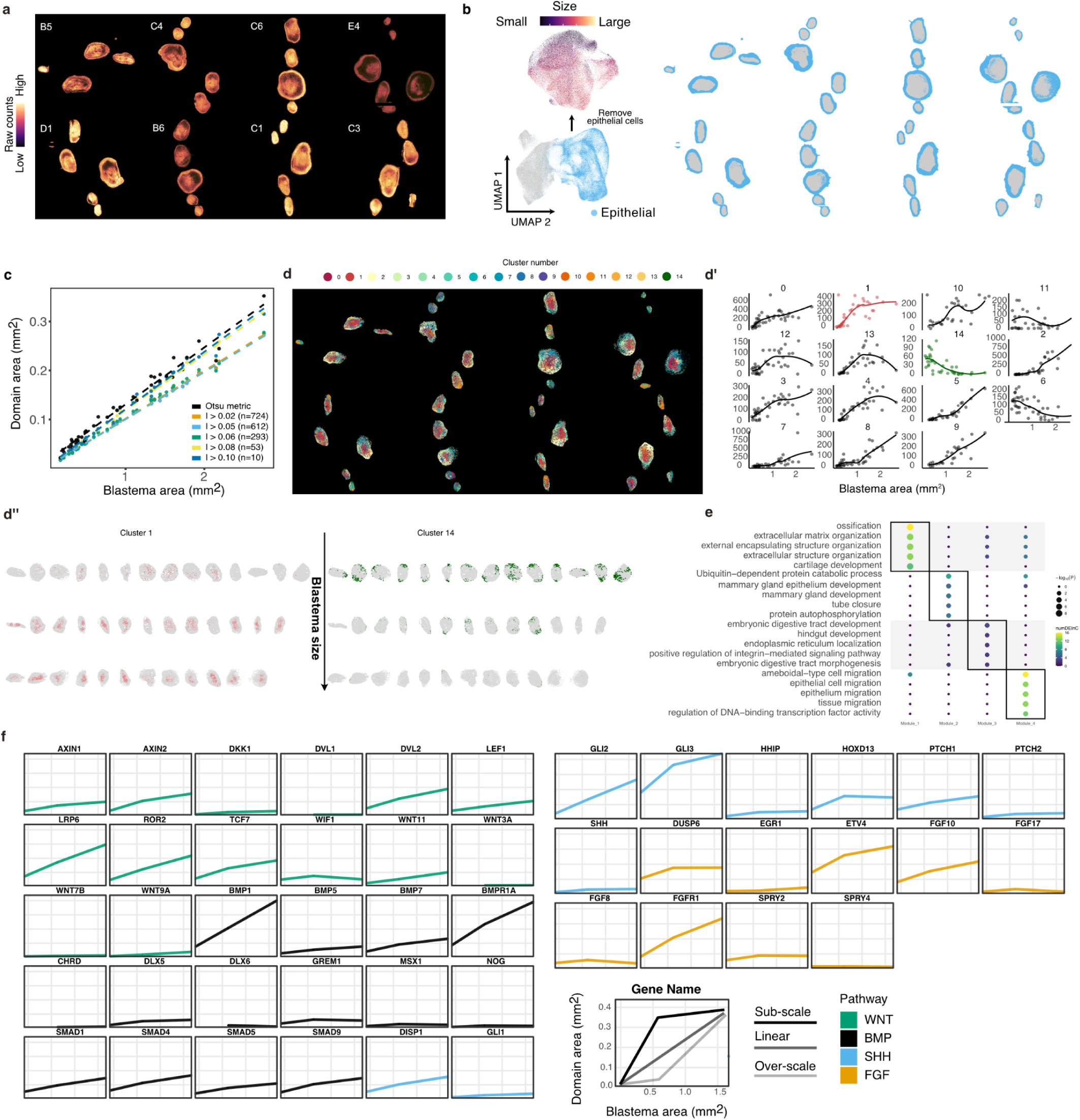
**a.** Total UMI counts across all batches. Epithelial regions display distinct count distributions compared to mesenchymal tissue.. **b.** Identification and removal of epithelial cells. Clustering (*Methods*) separates epithelial cells, shown in UMAP space and highlighted in blue in the spatial data. These cells are excluded from downstream analyses. **c.** Domain size scaling as a function of blastema area across different thresholds of the patterning index (n = number of genes), and using an alternative domain definition based on Otsu thresholding with the same gene set as in Fig. 1d. **d** Cluster size scaling for individual clusters. In **d′**, we highlight representative clusters that exhibit sub-scaling (1) or under-scaling behavior (14). In **d’’**, we show the spatial localization of cells belonging to these reference clusters. Blastema sizes are rescaled for visualization. **e.** Differential expression analysis and module identification of highly variable genes between small and large blastemas (Fig. 1f). GO enrichment analysis shows that genes downregulated in larger blastemas are associated with limb development related processes (right). Here, the top 1000 genes were used for the analysis, whereas 100 genes were used for visualization in Fig. 1f. **f.** Domain area as a function of the blastema area for representative genes in the highlighted pathways. Each point represents the average domain size across blastemas grouped into three size classes (small, medium, and large). The bottom right panel illustrates typical domain behaviors and their interpretation in terms of scaling. Domain and blastema area values are consistent across all panels and correspond to those shown in the schematic.

**Supplementary Figure 2:**
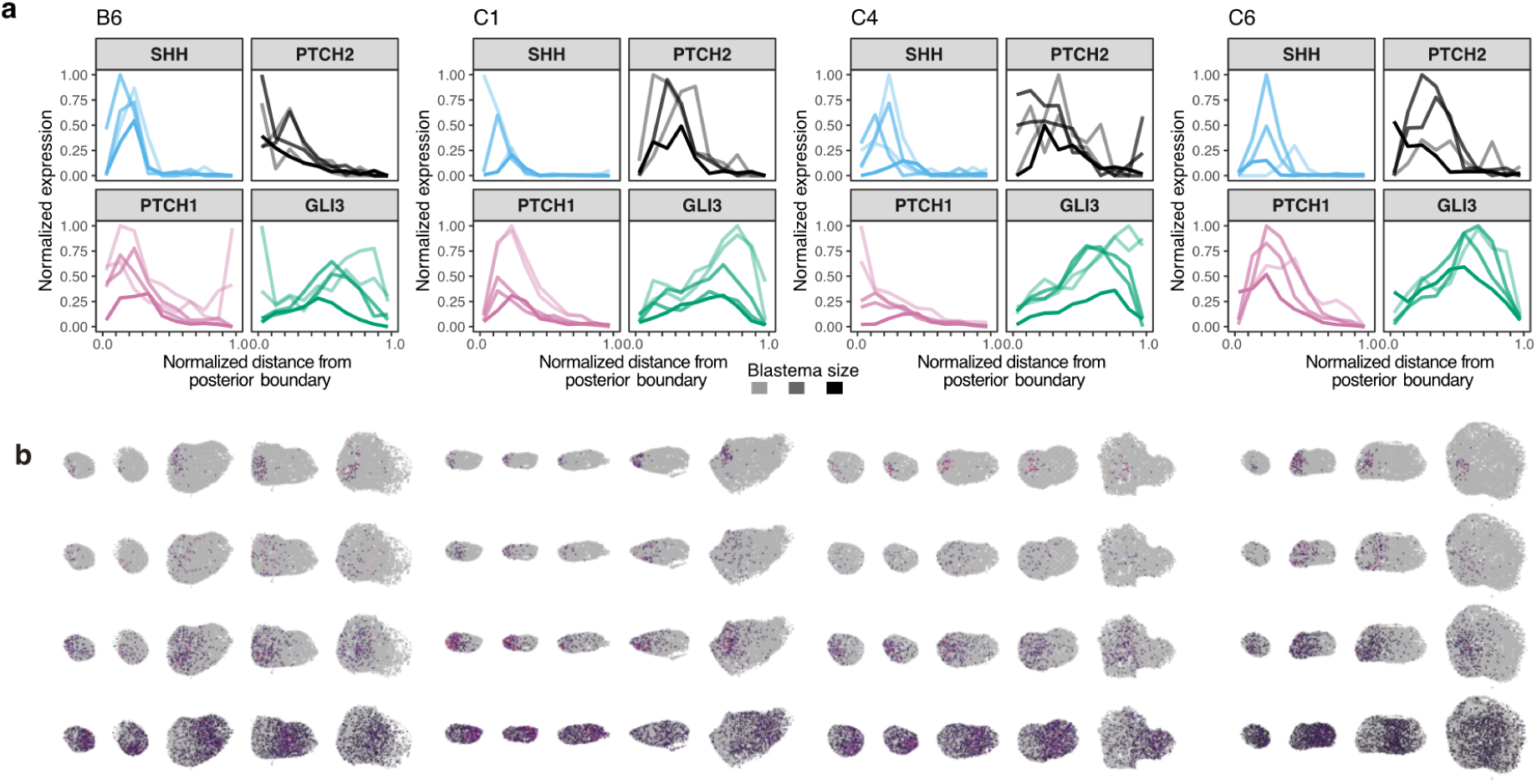
**a.** 1D profiles obtained for *Shh*, *Ptch1*, *Ptch2*, *Gli3* for three batches at early stage and one at a later stage (C1), showing reduced SHH signaling in the largest blastemas. **b**. Visualization of the normalised expression for the same genes across blastemas ordered by their sizes in each batch.

**Supplementary Figure 3:**
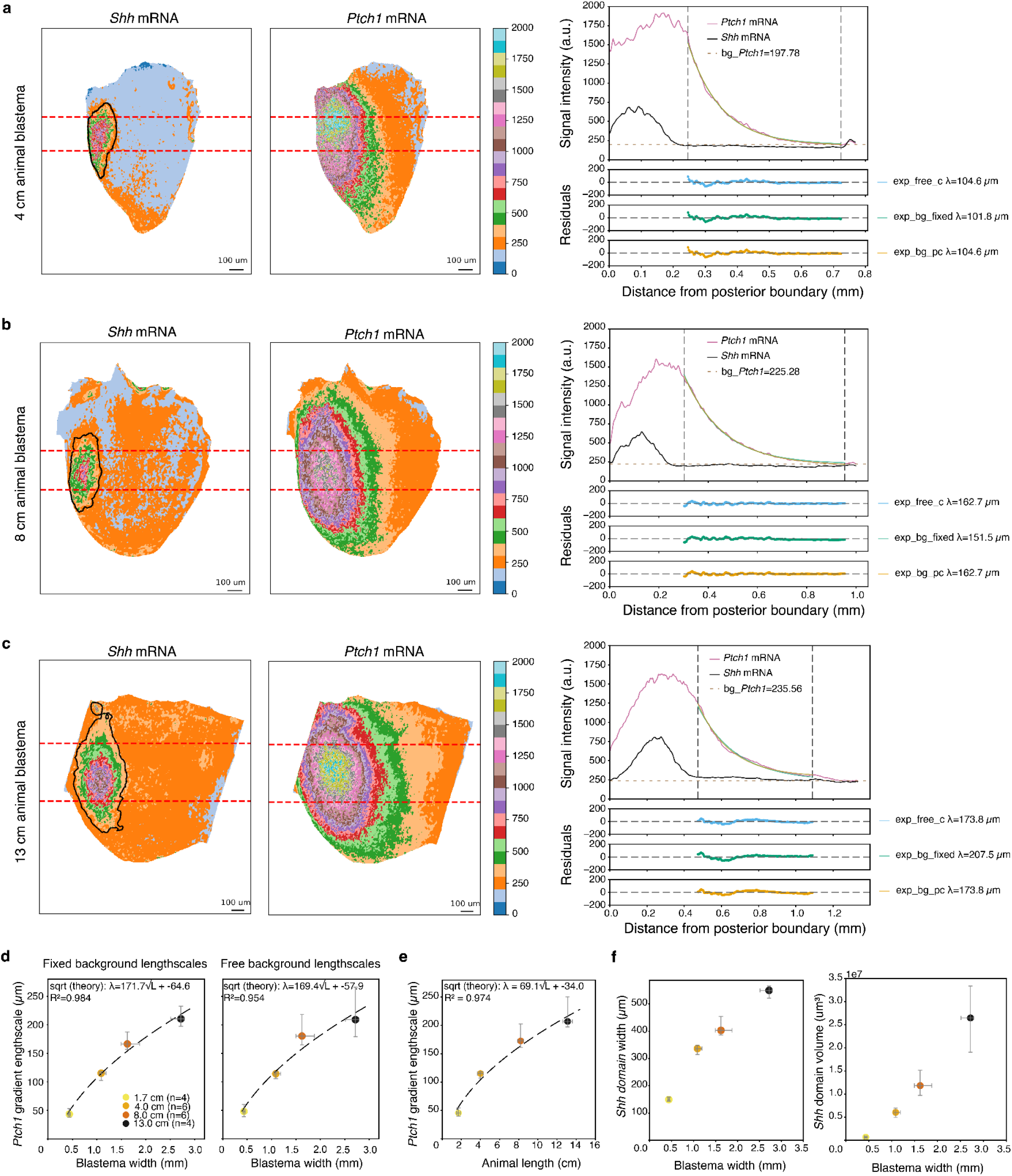
**a–c.** Representative light-sheet images and *Ptch1* gradient quantification for blastemas from 4 cm (a), 8 cm (b), and 13 cm (c) long animals. Left columns: *Shh* and *Ptch1* mRNA signal (HCR), average intensity projected, and colour-coded by intensity. Black outline, segmented *Shh* expression domain; red dashed lines, region used for intensity profile extraction. Scale bars, 100 µm. Right: *Ptch1* (magenta) and *Shh* (black) signal intensity profiles as a function of distance from the posterior boundary; dashed horizontal line, estimated *Ptch1* background. Vertical dashed lines indicate the exponential fit window. Below, residuals from three fit modalities: free intercept (blue, exp_free_c), fixed background (green, exp_bg_fixed), and partially constrained background (orange, exp_bg_pc), with fitted λ values indicated. Residuals are flat and near-zero across for all fits. **d.** *Ptch1* gradient lengthscale λ as a function of blastema width using fixed background (left) and free background (right) exponential fits. Points show median ± IQR. Dashed lines, square root fits confirming sublinear scaling is robust to fitting approach. **e.** *Ptch1* gradient lengthscale λ as a function of animal length (median ± IQR). Dashed line, square root fit, showing that sublinear scaling of the SHH gradient is also observed relative to animal size. **f.** *Shh* expression domain width (left) and volume (right) as a function of blastema width (median ± IQR), showing scaling of the *Shh* source with tissue size.

**Supplementary Figure 4:**
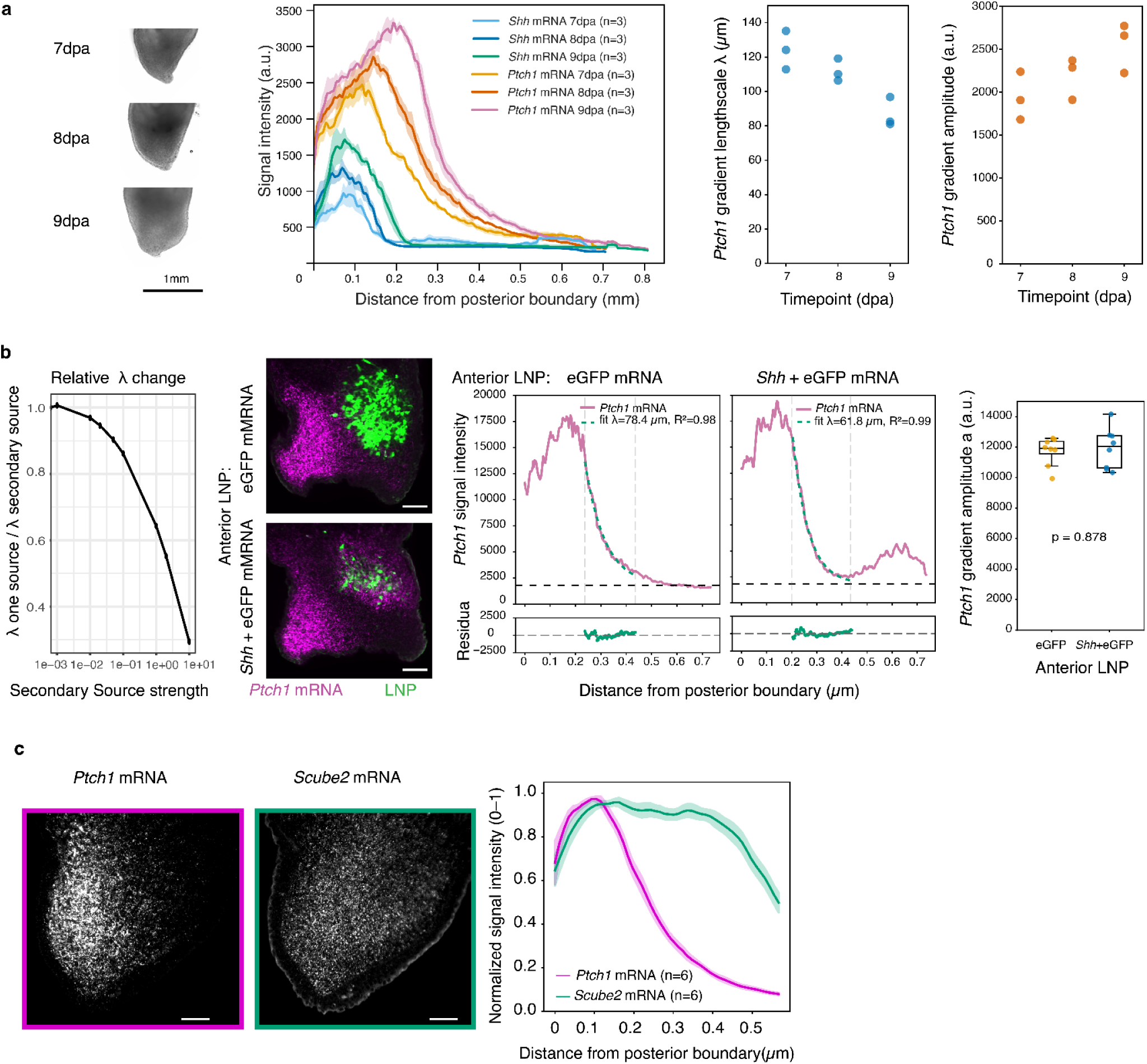
**a.** *Shh* and *Ptch1* mRNA expression profiles in regenerating blastemas (4 cm animals) at 7, 8, and 9 days post amputation (dpa). Left: representative widefield images of blastemas at each timepoint. Signal intensity (mean ± SEM) as a function of distance from the posterior boundary (n=3 per timepoint). As regeneration proceeds, *Ptch1* gradient amplitude increases and spatial range expands, while gradient lengthscale decreases, reflecting progressive sharpening and broadening of the SHH signaling domain. **b.** Left: Relative change in *Ptch1* gradient length-scale as a function of ectopic source strength, obtained from numerical simulations. The y-axis shows the ratio of the *Ptch1* gradient length-scale in the absence versus presence of an ectopic source of the strength indicated on the x-axis. Center-left: representative single-plane light-sheet images of blastemas injected anteriorly with eGFP LNPs (top) or *Shh* + eGFP LNPs (bottom). *Ptch1* mRNA HCR signal is shown in magenta, LNP distribution (eGFP expression) in green. Scale bar, 100 µm. Center-right:: Posterior-to-anterior *Ptch1* signal intensity profiles (pink) with exponential fits (dashed green lines) for anterior eGFP LNP, anterior *Shh* + eGFP LNP. Note that the fit window of eGFP controls is restricted to match the average fit-length of the Shh-injected condition. The dashed horizontal black line indicates the global background intensity estimated from control samples, used as a common intercept for all exponential fits. Right: Corresponding *Ptch1* gradient amplitudes are not significantly different between conditions (p = 0.878, Mann-Whitney U test), indicating that the observed length-scale reduction is not attributable to differences in overall signal amplitude. Boxes show median ± IQR; whiskers extend to 1.5× IQR; dots represent individual blastemas. n = 8 per group. **c.** Representative mRNA expression of *Ptch1* and *Scube2* in the axolotl limb blastema. Left, single-plane longitudinal optical sections showing *Ptch1* and *Scube2* mRNA (HCR in situ hybridization). Right, normalized signal intensity profiles along the anterior–posterior axis plotted as distance from the posterior boundary, showing broad *Scube2* expression relative to *Ptch1*. Lines show mean ± s.e.m.; *n* = 6 blastemas. Scale bars, 100 µm.

**Supplementary Figure 5:**
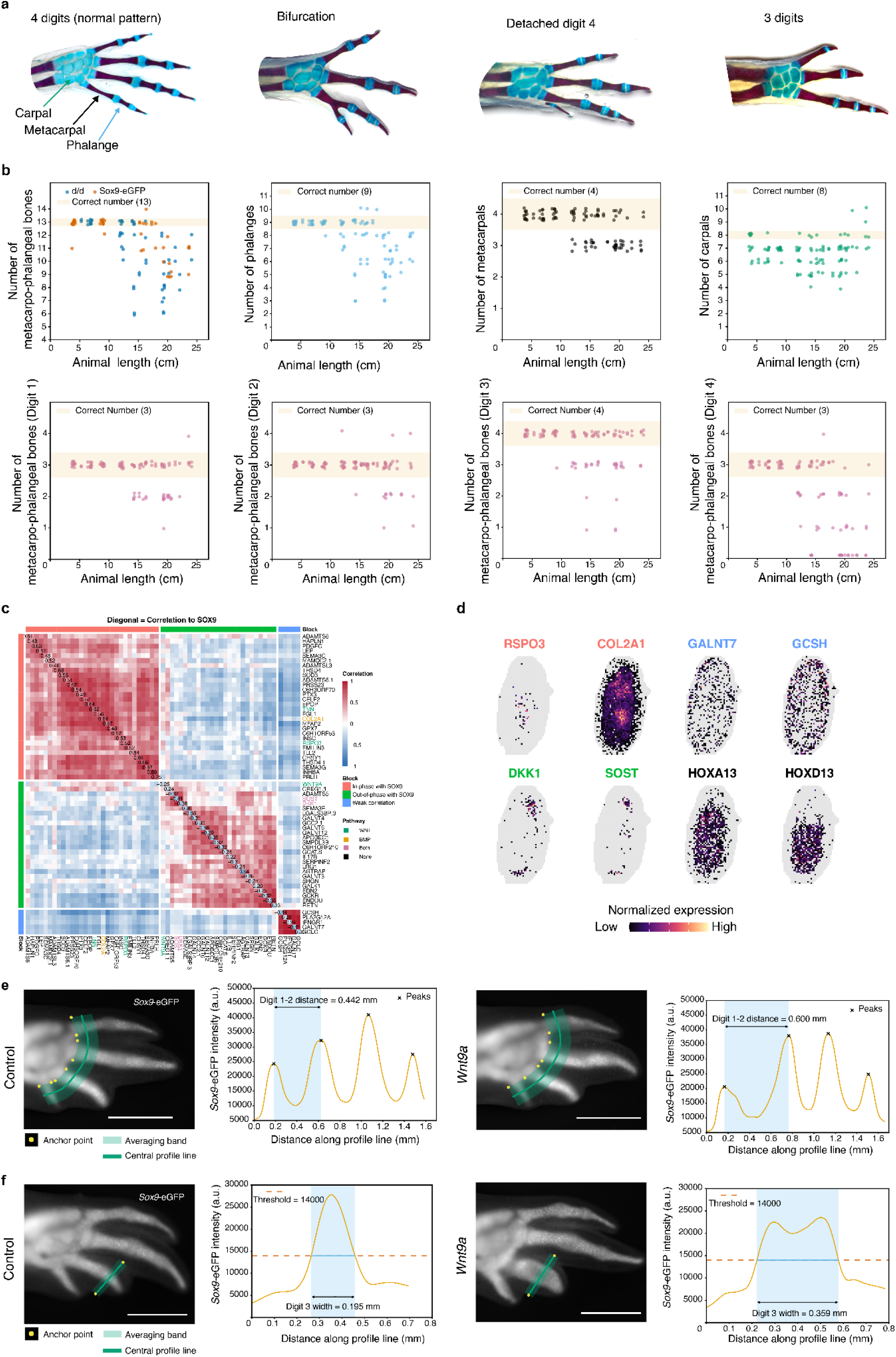
**a.** Representative Alcian Blue/Alizarin Red-stained limbs illustrating the digit scoring categories used in this study. From left: normal pattern (4 digits, with green arrow indicating carpals, black metacarpals, and pink phalanges); bifurcation, in which digit 3 bifurcates distally to produce a fourth digit tip; detached digit 4, in which a skeletal element is present but not connected to the carpal plate; and 3 digits. Bifurcation and detached digit 4 are both scored as 4 digits in quantifications but distinguished by “x” symbols in Fig. 3b. **b.** Individual skeletal element counts plotted against animal length. Top row, left: total metacarpo-phalangeal bone count in d/d (blue) and *Sox9-eGFP* (orange) animals; beige shading indicates the correct element count (13). Top row, centre-right: number of phalanges, metacarpals, and carpals, with correct element counts indicated (9, 4, and 8 respectively). Bottom row: phalangeal counts per digit (digits 1-4), with correct phalangeal numbers indicated (2, 2, 3, and 2 respectively). Beige shading indicates the correct element count in all panels. Reduced regeneration fidelity with increasing animal size is observed for phalanges, metacarpals, and per-digit phalangeal counts, and in both genotypes; carpal counts are reduced across all animal sizes. Each point represents an individual limb (n=117 limbs). **c.** Correlation matrix obtained by projection of the two-dimensional UMI counts from the medium blastema of the C1 batch along a one-dimensional line orthogonal to the SOX9 profile. On the diagonal we show the correlation of each of the most highly differentially expressed genes in the chosen blastema to SOX9. We indicate with different colours if the chosen genes are in WNT/BMP/both pathways. The matrix is organised into three blocks of which one-dimensional schematics are shown on the right (black) and compared to SOX9 (purple) **d.** Representative spatial expression patterns from the same blastema as in c. Two genes are shown for each correlation block: *Rspo3* and *Col2a1* (in phase with *Sox9*), *Dkk1* and *Sost* (out of phase), *Galnt7* and *Gcsh* (weak correlation) as well as distal Hox genes *Hoxa13* and *Hoxd13*. **e.** Quantification of digit 1-2 spacing in representative *Sox9*-eGFP limbs corresponding to Fig. 3e. Left: Control limb (widefield image) with thick spline profile line (fixed-width averaging band) drawn from manually placed anchor points at the base of the digits, and corresponding *Sox9*-eGFP signal intensity projected onto the central profile line. Right: WNT9a LNP-injected limb and corresponding profile. Peaks are automatically detected; digit 1-2 distance is extracted as the distance between the first two peaks. Scale bar, 1 mm. **f.** Quantification of digit 3 width in representative *Sox9*-eGFP limbs corresponding to Fig. 3f. *Left:* Control limb (widefield image) with thick profile line (fixed-width averaging band) drawn between manually specified anchor points across digit 3 at the level of the injection site (see main figure) and corresponding *Sox9*-eGFP signal intensity projected onto the central profile line. *Right:* WNT9a LNP injected limb and corresponding profile. Digit 3 width is extracted as the peak width at a fixed intensity threshold applied uniformly across all samples (dashed line, threshold = 14000 a.u.). Scale bar, 1 mm.

**Supplementary Figure 6:**
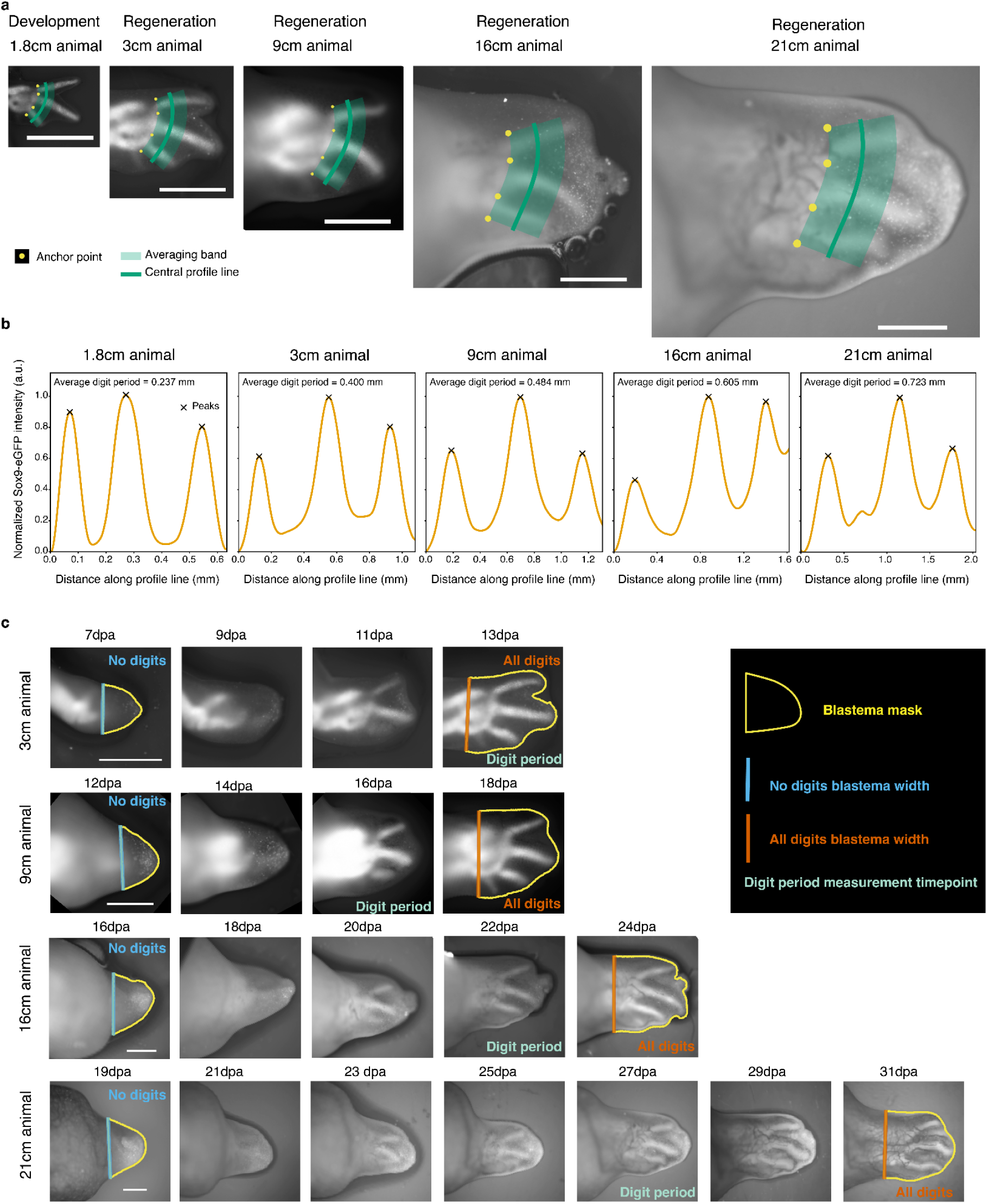
**a.** Representative *Sox9*-eGFP images of developing (1.8 cm) and regenerating (3, 9, 16, 21 cm) limbs at the timepoint of digit period extraction (when 3 digits are clearly separated). Yellow dots, manually placed anchor points along the base of the digits; green shaded band, averaging band extending distally to the length of digit 3 and perpendicular to the digit rays; dark green line, central profile line onto which intensities across the band width are averaged. Scale bars, 1 mm. **b**. Normalized *Sox9*-eGFP intensity profiles along the central profile line for each limb shown in a. Profiles were smoothed with a Savitzky–Golay filter; crosses mark detected peaks corresponding to digit positions. Average digit period (mean inter-peak distance between digits 1-2 and 2-3) is indicated for each limb. **c**. Time-course *Sox9*-eGFP live imaging of regenerating limbs across animal sizes (3, 9, 16, and 21 cm). Yellow outline, blastema mask; blue line, blastema width at the "no digits" stage (before digit emergence), measured at the base of the blastema; orange line, blastema width at the "all digits" stage (when the complete digit series is first present), measured at the base of the hand field, defined as the proximal boundary of *Sox9* expression at the level of the forming carpals (see Methods). "Digit period" labels indicate the timepoint used for digit periodicity measurements (see a, b). For limbs that failed to produce a fourth digit, the "all digits" stage was defined as 4 days after the digit period measurement timepoint. A detached *Sox9*-positive element was considered equivalent to a fourth digit for stage assignment. Scale bars, 1 mm. dpa, days post amputation.

**Supplementary Figure 7:**
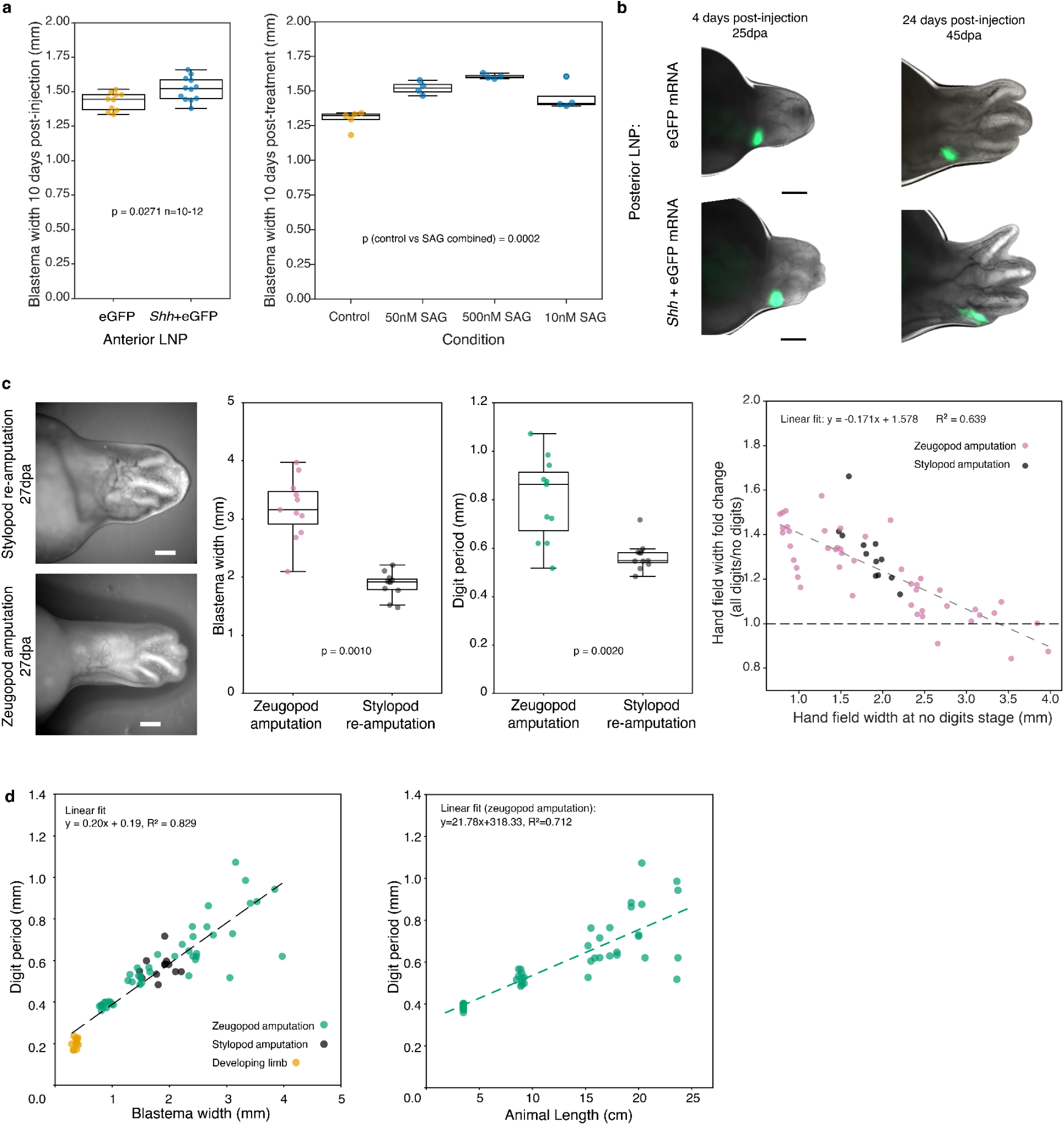
**a.** Left: anterior injection of Shh + eGFP mRNA LNP increases blastema width (assessed at 17dpa, 10 days after injection) compared to eGFP control. Right: SAG treatment increases blastema width (assessed at 17dpa, 10 days after initiation of a 2 days treatment) **b.** Posterior LNP injection in a large axolotl blastema. Top row, eGFP mRNA control LNP; bottom row, *Shh* + eGFP mRNA LNP. Left, widefield images at 4 days post-injection (25dpa) showing LNP localisation in the posterior blastema. Right, images at 24 days post-injection (45dpa). Scale bars, 1 mm. **c.** Left: representative widefield images of the same limb at 27dpa following zeugopod amputation (top) and stylopod re-amputation (bottom), showing a visibly smaller blastema and restored digit number after re-amputation. Scale bars, 1 mm. Center-left, blastema width is significantly reduced following stylopod re-amputation compared to zeugopod amputation (paired Wilcoxon signed-rank test, P = 9.77 × 10⁻⁴; n = 11 limb pairs). Center-right, digit period is correspondingly reduced (P = 1.95 × 10⁻³). Right, blastema width fold change (all digits / no digits stage) plotted against initial blastema field width for all zeugopod amputations (replotted from Fig. 4c) and stylopod re-amputations (black). Stylopod re-amputation blastemas fall on the same size-dependent growth trend as zeugopod blastemas(R² = 0.639; y = −0.171x + 1.578). Dashed line at fold change = 1 indicates no expansion. Box plots show median, interquartile range, and whiskers extending to 1.5× IQR. **d.** Left, digit period scales linearly with blastema width (measured before digit emergence) across regenerating limbs from zeugopod amputation (teal), stylopod re-amputation (black), and developing limbs (orange). n= 69 limbs. Right, digit period plotted against animal length (zeugopod data only. n= 45 limbs.

