## Supplementary Theory for "Adult regenerative defects arise from discordant scaling of signal dependent growth and patterning"

#### Introduction

In this Supplementary material we give details in support of the theory described in the main text. In Sec. 1 we introduce the set of minimal equations describing scaling of SHH pathway via an expansion-repression mechanisms. In Sec. 2 we provide details on the analytical and numerical solutions to the equations derived in the previous section. In Sec. 3 we discuss alternative mechanisms of scaling proposed in literature. Finally, in Sec. 4, we show a mechanisms for the scaling of digit periodicity and digit loss.

#### 1 Model definition

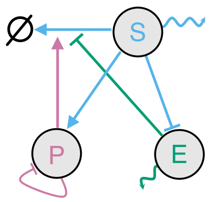


Figure ST 1: Minimal model describing the SHH-Ptch1 interaction network (resp. S and P), and its coupling to an expansion-repression mechanism (expander E). Detailed interactions are given in Eq. (2).

Here we consider a minimal model for the establishment of an exponential SHH gradient which scales with the system size. As we are interested in one-dimensional gradients along their AP axis, we will restrict our theory to a one-dimensional domain and discuss later the effect of two and three spatial dimensions. We restrict ourselves to the simplest regulatory network able to recapitulate the experimental findings and which is able to predict perturbations. To this end, we consider only three genes, SHH ($s$) , PTCH1 ($p$) and the expander ($e$) which interactions are shown in Fig. ST 1. For each gene, we first derive the equations both for the mRNA levels (subscript m), which is the quantity actually observed in spatial transcriptomics, as well as the protein concentration (substract P), which is the quantity which diffuses spatially and feeds back onto other species.

Based on the experimental findings (Fig. 2), the production of Shh mRNA ($s_m$) is localized in space, so that we can take a simple mathematical form, $\alpha_s\Theta(yL - x)$, where $y$ is the fraction of the source with respect to the total system size $L$ and $\alpha_s$ is the production rate. SHH proteins diffuse with a rate $D_s$ and based on previous findings [1, 2] they are degraded as a complex with its receptor PTCH1 at a rate $k_{ps}$, as well as with a baseline rate $k_{se}$. Conversely, PTCH1 protein ($p_p$) do not diffuse and are degraded with SHH at rate $k_{ps}$, as well as with a baseline rate $k_{pP}$.

They are produced proportional to the level of Ptch1 mRNAs, which are themselves produced at rates which increase with increasing SHH levels. To this end, and based on previous findings [1], we model the production of Ptch1 mRNA via a Hill-type functions with rate $\alpha_p$ and Hill-coefficient $H_p$, which depends positively on SHH protein levels and negatively on PTCH1 protein levels (as in the absence of SHH, PTCH1 would effectively self-repress). It is possible to make different choices for this function; however, we found this to be a choice consistent with the experimental findings.

In addition to these regulatory interactions within the SHH pathway we introduce a third specie, the expander ($e$) [3, 4], which is able to modulate the spatial profile of SHH. We have mainly two possibilities to introduce such a molecule into the equation: it can increase the diffusion, or decrease the degradation, of the morphogen. Here, we consider that the expander prevents SHH degradation, in 4 we justify the choice between degradation and diffusion. For the dynamics of the expander mRNA $e_m$, as it is unknown, we consider the minimal function as in [3], where the rate of the production $\alpha_e$ is inhibited by SHH with a Hill-type relationships with coefficient $H_e$ and saturation $TT$. The expander protein $e_P$ degrades with a constant rate $k_e$ and diffuses with a rate $D_e$. All together the system is described by the following set of differential equations,

$$
\begin{aligned}
s_m &= \alpha_{sm}\Theta(yL - x) \\
\partial_t s_P &= \alpha_{sT} s_m - k_{ps\,P} p_P s_P - k_{se\,P}\frac{s_P}{1+e_P} + D_s \partial_{xx} s_P \\
\partial_t p_m &= \alpha_{pm}\frac{s_P^{H_p}}{p_P^{H_p} + s_P^{H_p}} - k_{p\,m} p_m \\
\partial_t p_P &= \alpha_{pT} p_m - k_{ps\,P} p_P s_P - k_{pP} p_P \\
\partial_t e_m &= \alpha_{em}\frac{TT^{H_e}}{TT^{H_e} + s_P^{H_e}} - k_{em} e_m \\
\partial_t e_P &= \alpha_{e\,T} e_m - k_{eP} e + D_e \partial_{xx} e_P
\end{aligned}
\tag{1}
$$

The previous equations, supported with boundary and initial conditions, fully defines the spatio-temporal dynamics of the SHH, PTCH1 and the expander proteins and mRNA. Numerical solutions of the previous equations can be directly compared to experimental results were only mRNA was measured. However, in order to understand the origin of scaling and several features of morphogen patterns we integrate out the equation for mRNA by taking the limit of fast translation. We obtain the more compact equations for the proteins (removing the superscript P and redefining the parameters),

$$
\begin{aligned}
\partial_t s &= \alpha_s \Theta(yL - x) - k_{ps} p s - k_{se}\frac{s}{1+e} + D_s \partial_{xx} s \\
\partial_t p &= \alpha_p \frac{s^{H_p}}{p^{H_p} + s^{H_p}} - k_{ps} p s - k_p p \\
\partial_t e &= \alpha_e \frac{TT^{H_e}}{TT^{H_e} + s^{H_e}} - k_e e + D_e \partial_{xx} e
\end{aligned}
\tag{2}
$$

If not specified otherwise, the parameters chosen for the simulations for the SHH and PTCH1 follow closely the one in [1] and are provided in Table 1. Numerical simulations are performed with a 2nd order Runge-Kutta integration in time and finite difference in space with no-flux boundary conditions.

| Model parameter | Description | Value |
|---|---|---|
| $\alpha_s$ | Source strength of $s$ | $10^{-1}$ |
| $k_{ps}$ | Degradation rate of $p$ as a complex with $s$ | $10^{-4}$ |
| $k_{se}$ | Degradation of $s$ | $10^{-1}$ |
| $D_s$ | Diffusion coefficient of $s$ | 10 |
| $\alpha_p$ | Production rate of $p$ | 1 |
| $k_p$ | Degradation rate of $p$ | $10^{-1}$ |
| $H_p$ | Hill coefficient for $p$ activation | 5 |
| $\alpha_e$ | Production rate of expander $e$ | $10^{-2}$ |
| $k_e$ | Degradation rate of expander $e$ | $10^{-4}$ |
| $D_e$ | Diffusion coefficient of expander $e$ | 10 |
| $H_e$ | Hill coefficient for expander regulation | 5 |
| $y$ | Relative size of source region | $L/4$ |

Table 1: Parameters used in the minimal PDE model. All parameters are taken in dimensionless units.

#### 2 Scaling of Morphogen gradients

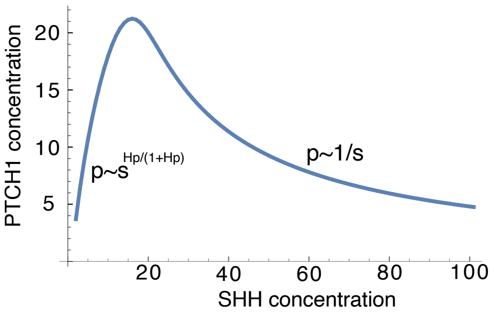


Figure ST 2: PTCH1 concentration as a function of SHH concentration, where we highlight the relationship at low and high concentration of SHH.

In this section we provide details of the analytical derivation of the shape of morphogen and expander profiles, which we confirm numerically and test against experimental evidences. To get analytical insight we consider an expander profile almost constant along the tissue, which is valid for $D_e \gg 1$ such that we replace, $e \to \langle e \rangle_x / L = \bar{e}$. In this limit, the solution for Ptch1 level at steady state is

$$p = \alpha_p \frac{s^{H_p}}{p^{H_p} + s^{H_p}} \frac{1}{k_{ps} s + k_p} \tag{3}$$

which we show in Fig. ST 2. In order to get more insights, we find the following scaling limits:

$$\begin{aligned} s \to 0\,, \quad & p \sim s^{\frac{H_p}{H_p+1}} \\ s \to \infty\,, \quad & p \sim 1/s \end{aligned} \tag{4}$$

Already from the steady-state solution we have two non-trivial observations. Firstly, for low values of SHH, PTCH1 is directly proportional to SHH, meaning that the tails of the SHH morphogen gradient and its target, PTCH1, have the same functional form, such that if SHH is an exponential, the same holds for PTCH1. We then need to solve for only one of them. Secondly, for high concentration of SHH, PTCH1 is inversely proportional to SHH. We can conclude that first, a decrease of SHH lead to an increase of PTCH1 and second, after a typical length-scale, they decay proportionally. Following this arguments, there is a peak of PTCH1 which can lay outside the SHH domain and that is what we observed in the experiments in Fig. 2 of the main text. We then chose parameters such that there exist a peak of PTCH1 outside the SHH domain.

As we explained below, far from the source, PTCH1 and SHH decay with the same functional form. We take now the limit $k_{ps} \ll k_{pe}$ (justified for the existence of a PTCH1 peak), such that $p$ is treated as a small perturbation. In this limit, the stationary equation for $s$ is solved by an exponential function with a length scale $\lambda_s = \sqrt{\frac{D_s(1+\bar{e})}{k_{se}}}$. Thus, with the previous scaling $p$ will initially increase for large value of $s$ and will then decay exponentially with a length scale $\lambda_p = \lambda_s / \frac{H_p}{H_p+1}$. This finding implies that PTCH1 has a peak which is shifted compared to the SHH domain. In Figure S3 of the main text, we observe such shift across several animal sizes. All together for $x \to \infty$

$$\begin{aligned} s &\sim \exp\left[-x/\sqrt{D_s(1+\bar{e})/k_{se}}\right] \\ p &\sim \exp\left[-x/\sqrt{D_s(1+\bar{e})(H_p+1)^2/k_{se}H_p^2}\right] \end{aligned} \tag{5}$$

In Fig. ST 3 we show the numerical versus the analytical prediction of the scaling of SHH for two systems with a size smaller (left) and larger (right) than the typical length-scale of the expander, $\lambda_e = \sqrt{D_e/k_e}$ .

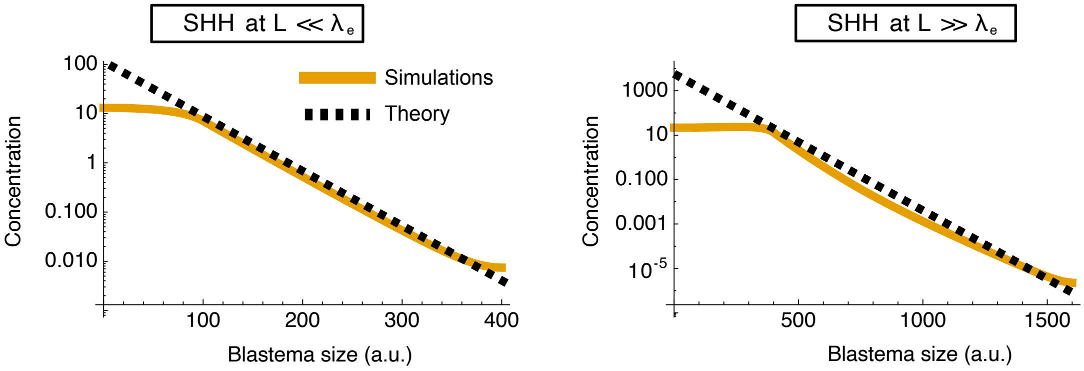


Figure ST 3: Numerical versus analytical predictions of the SHH profile from Eq. (5) when the system size is greater (left) or smaller (right) than the typical length-scale of the expander.

#### 2.1 Scaling solution for the expander profile

In order to find the value of $\bar{e}$ to close the system of equation, we need to solve at steady state the following equation,

$$-D_e\,\partial_{xx}e + k_e\,e \;=\; \alpha_e\,\frac{TT^{H_e}}{TT^{H_e}+s^{H_e}(x)}, \qquad e'(0)=e'(L)=0. \tag{6}$$

Here we solve the equations self-consistently, meaning that we look for a solution $s(x)$ which is system size dependent and check *a posteriori* that such a solution satisfies Eq. (6). As a first approximation (which will be sufficient) we replace the expander Hill function by

$$\frac{TT^{H_e}}{TT^{H_e}+s^{H_e}(x)} \;\approx\; \begin{cases} f_0, & 0 \le x \le a, \\ 1, & a < x \le L\,, \end{cases} \tag{7}$$

where $f_0$ and $a$ will be determined self-consistently by imposing the values of the expander at the boundaries. and $q = \alpha_e/k_e$. In this approximation, Eq. (6) is straightforwardly solved by

$$e(x; L) \approx \begin{cases} q\Big[f_0 + A(L)\cosh\big(\frac{x}{\lambda_e}\big)\Big], & 0 \le x \le a, \\ q\Big[1 - B(L)\cosh\big(\frac{L-x}{\lambda_e}\big)\Big], & a \le x \le L, \end{cases} \tag{8}$$

with

$$A(L) = \frac{1 - f_0}{\cosh\left(\frac{a}{\lambda_e}\right) + \sinh\left(\frac{a}{\lambda_e}\right)\coth\left(\frac{L-a}{\lambda_e}\right)}, \qquad B(L) = A(L)\,\frac{\sinh\left(\frac{a}{\lambda_e}\right)}{\sinh\left(\frac{L-a}{\lambda_e}\right)}. \tag{9}$$

Let us assume that $e(0) = e_0$ and $e(L) = e_L$ are known, upon introducing the dimensionless variable $\Lambda = \frac{L}{\lambda_e}, \alpha = \frac{a}{\lambda_e}$ and the rescaled boundary values $u = \frac{e_0}{q}, \quad v = \frac{e_L}{q}$ we evaluate (8) at the boundaries and with the continuity conditions at $x = a$ one finds the simple relations

$$u = f_0 + (1 - f_0)\,\frac{\sinh(\Lambda - \alpha)}{\sinh\Lambda}, \qquad v = 1 - (1 - f_0)\,\frac{\sinh\alpha}{\sinh\Lambda}. \tag{10}$$

When can then eliminate $f_0$ and have a closed form for $\alpha$ , then, provided $|t| < 1$,

$$\alpha = 2\ \mathrm{artanh}\left(\frac{\cosh\Lambda - \frac{u-1}{v-1}}{\sinh\Lambda}\right), \qquad a = \lambda_e\,\alpha \tag{11}$$

and $f_0$ follows from either identity in (10), e.g.

$$f_0 = \frac{u\sinh\Lambda - \sinh(\Lambda - \alpha)}{\sinh\Lambda - \sinh(\Lambda - \alpha)} \tag{12}$$

These formulas are valid when $0 < \alpha < \Lambda$, $0 < f_0 \le 1$. In Fig. ST 4 we show the comparison of the analytical results with numerical simulations in the cases when the typical length-scale of the expander, $\lambda_e$, is greater or shorter than the system-size.

We now seek to understand the dependence of the length scale with the expression of SHH. To this end, we fix the system size and vary the parameter $\alpha_s$. The results are shown in Fig. ST 6, where we observe that an increase of the expression of SHH decreases the length-scale of Ptch1 as observed in experimental measurement.

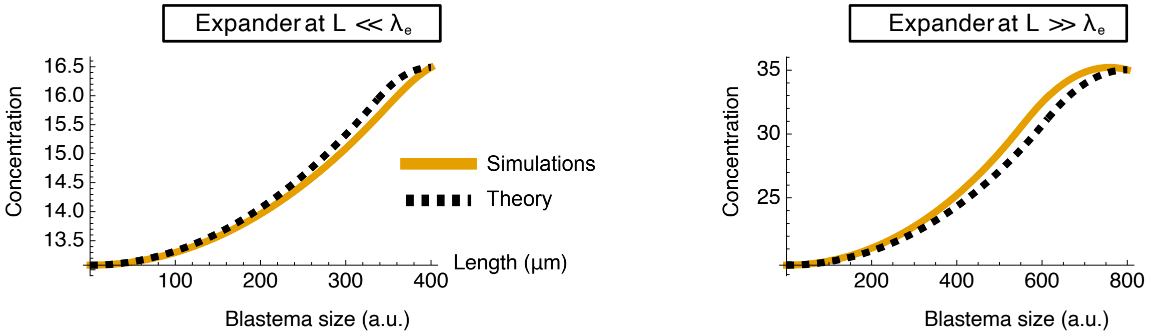


Figure ST 4: Numerical versus analytical predictions of the expander profile when the system size is greater than the typical length-scale of the expander (left) or the opposite (right). For small system sizes, the expander achieves almost a uniform profile.

#### 2.2 Limits of scaling

Our model in Eq. (2) does not have any exact solution. In general we expect two scenarios which are dependent on the values of the parameters. First, a good approximation of the length scale of the gradient is a scaling with system size as $\lambda_s \sim L^{1/2}$, consistent with the scaling of transcriptionally similar clusters in Fig. 1e. Alternatively, we could observe a linear increase of the length scale of the gradient followed by a saturation at large system sizes. We define this regime as sub-scaling in the main text. Irrespective of the specific scaling relationship, the system cannot sub-scale indefinitely (Figure 2d) and in this section we prove that there is a limit size above which the length-scale is saturated as observed for example in *ex vivo* models of embryonic anterior-posterior patterning [5]. Upon studying the limits of Eq. (2), we find that even in case of very fast diffusion of the expander $D_e \gg D_s$, there is a maximal length scale. In particular the equation for the expander, gives that the maximal value for the expander concentration at the tail of SHH, is $e^* = \alpha_e/k_e \gg 1$. Putting this result back in the equation for $s$, as in the tail the term $k_{ps}ps$ is negligible, we find that $s$ has a maximal length-scale $\lambda_s^* \approx \sqrt{\frac{D_s \alpha_e}{k_e k_s}}$ and similarly for PTCH1, $\lambda_p^* \approx \sqrt{\frac{D_s \alpha_e (H_p+1)^2}{k_e k_s H_p}}$. In Fig. ST 5 we show the provided maximal length-scale in comparison with the scaling of the pattern length-scale over several decades of system size.

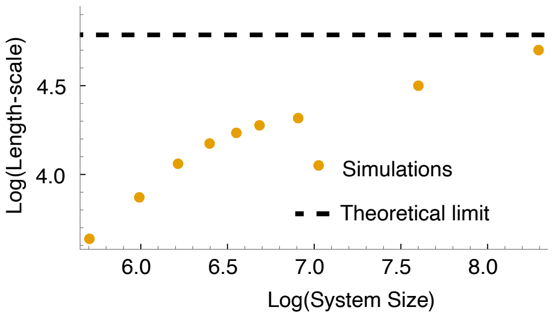


Figure ST 5: Scaling of the Ptch1 gradient length-scale to the system size, the black dashed line denotes the maximal length-scale and so the limit of scaling.

#### 2.3 Prediction of experimental perturbations

The length-scale of the SHH and PTCH1 gradients are not only determined by the system size via the expander concentration, but they can be affected by other factors, such as the size of the source or its amplitude. In particular we observe that an increase in the SHH expression in time lead to a slight decrease of the length-scale of the pattern. To this end, we numerically simulate equations (2) by keeping all the parameters fixed and varying only the amplitude of the source $\alpha_s$. In Fig. ST 6 we show the results of these simulations with the maximal and minimal length-scale of the system. It is evident that the length-scale of the gradient decreases with an increasing amplitude of the source. Intuitively, this phenomena is caused by the size of the system. Specifically, for a system much larger than the typical length-scale of the pattern we do not observe any deviation. However, when the space is limited, due to reflecting proximal boundary conditions, the gradient can not expand throughout the tissue, thus an increase in amplitude leads to a decrease of the length-scale.

### 3 Different scaling hypothesis

#### 3.1 Growth

One possibility for scaling of morphogen gradients is coupling patterning with tissue growth. The mechanisms underling scaling is driven by advection-dilution terms due to proliferation which increase in time [6, 7]. Despite this mechanisms has been proposed in other context, such as development of the *Drosophila* wing, it cannot be the main cause of scaling during axolotl limb regeneration. First, we observe scaling not in time but between animals of different sizes. Moreover, the gradient is established in the amputated stump even before the morphological appearance of the blastema. If it would be due to growth, than we should compare the time-scale of growth between smaller to larger individuals, which are much larger than typical protein half-life time-scale. Second, in (Fig. S4 in the main text) we observe during growth a shortening of the length-scale of the gradient. This is probably due to boundary effects, but it is again another indication that growth is not affecting the scaling of the gradient. Third, in (Fig. 2h in the main text) we found that an additional endogenous source of SHH decreases the length-scale of the gradient, which would lead to the opposite scenario if growth is the cause of the gradient expansions. Taken together, despite we cannot exclude that growth changes the amplitude, we do not have any evidence that points toward an effect on the scaling of the gradient.

#### 3.2 Recycling

In the context of DPP gradient, recycling has been shown to play a critical role during scaling of the gradient [8, 9]. However, despite recycling might lead to a change of the expander profile, in the known recycling models an expander-like mechanisms or a coupling to growth is required to scale the length-scale of the gradient. Altogether, we can not exclude that recycling is involved in the formation of the SHH gradient as well as playing an important role in how it affects the length-scale. Our minimal model is intended to show the failure of scaling and a size-sensing mechanisms, not the precise biochemical mechanisms.

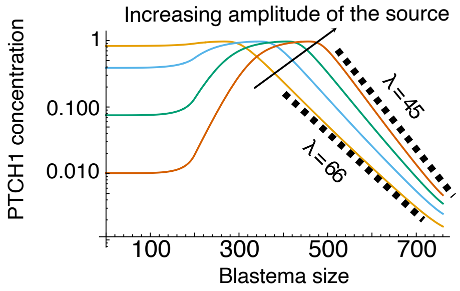


Figure ST 6: Scaling of Ptch1 with increase production of SHH. The length-scale of the pattern is inversely proportional to the increase of SHH expression

#### 3.3 Mechanical forces

Finally, we note that there has been some alternative proposals compared to reaction-diffusion to understand spontaneous patterning, in particular based on mechanical forces. Murray-Oster mechano-chemical models, for instance, were specifically designed with chondrogenesis and cartilage condensation in mind, proposing that positive feedbacks between cellular traction forces, ECM stresses and cell density could drive periodic condensation akin to digit formation [10, 11]. Such a mechanism has been shown to operate in settings such as follicle formation in chick [12], and mechanical forces have indeed been recently implicated in digit formation in mice [13]. Theoretically, it has also been shown that condensation could modulate morphogen reaction-diffusion by creating advective terms that create qualitatively different types of instabilities [14]. Yet, we note that in all of these cases, the length scale of periodic digit patterning is not expected to scale with field size *per se*, and additional mechanisms still have to be invoked. Furthermore, for any type of mechanical instability, we will expect that digit number will arise from a ratio of intrinsic patterning length scale and field size, meaning that the scaling arguments we make for Fig. 3 are expected to hold, although exploring the explicit potential role for mechanics in setting up or modulating the length scale of patterning would be an interesting next step.

### 4 Turing-like mechanism predicts digit patterning

In this section, we derive a minimal model to explain how larger animals often fail to regenerate the correct number of fingers (Fig. 3a). From the spatial transcriptomic (Fig. 1f) and microscopy data (Fig. 3a), we found that the pathways which are activated during digit regeneration (WNT and BMP) typically scale better than the SHH pathways. Specifically, we found evidence that the typical length-scale set by the digit period scale linearly with the system size (Fig. 4b). Interestingly, typical patterning mechanisms, such as the Turing model have a length-scale which is given by the microscopic molecular mechanisms and thus is not size-dependent. As we are going to show later, size-dependent parameters can allow Turing patterns to scale, but it would give rise to scaling of the entire field, such that it is not possible to justify within this model why there are less digits. Here we do not comment on the reason of the scaling of the BMP-WNT pathways, rather we try to understand the reason why in larger animals a digit is often missing. The simplest explanation would be that the loss of the last digit is related to the loss of scaling of digit periodicity. However, this would not result in both a scaling of the BMP and WNT pathway and in that case, the animal could actually develop more digits rather than less, Section 4.2. We then reject such hypothesis and look for a different minimal mechanism. It was already shown in [15] that SHH is responsible for the growth of the limb, such that a total inhibition would result in a disappearance of three out of four finger and a blastema which did not growth in size. To this end, we make the simple relationship, that whatever the mechanism for the formation of the fingers, if the growth of the blastema fails for larger animals, due to the failure of scaling of SHH, than we expect a feedback into the mechanisms of finger formation. As the mechanism for the formation of fingers we choose a Turing system, based on previous findings in mouse [16] and our perturbation experiments, but we emphasize that the mechanism that we propose does not depend on the specific implementation of the patterning mechanism, but rather on the differential scaling of patterning vs growth.

The Turing model that we consider is a three node gene network, BMP,WNT and SOX9 where BMP and WNT act respectively as activator and inhibitor of SOX9. In [16] it was considered that both WNT and BMP are self-inhibiting, however as at linear level degradation and self-inhibition cannot be distiguished, we simply the network interpretation by considering only degradation. This leaves at the level of linear stability analysis the Turing network of [16] invariant. The equations describing the dynamics of perturbation around an homogeneous solutions are given by,

$$\partial_t sox(x,t) = \alpha_{\rm s} + k_2\, bmp(x,t) - k_3\, wnt(x,t) - \big(sox(x,t) - sox_0\big)^3 \tag{13}$$

$$\partial_t bmp(x,t) = \alpha_{\rm b} - k_4\, sox(x,t) - k_5\, bmp(x,t) + D_b\, \partial_{xx} bmp(x,t) \tag{14}$$

$$\partial_t wnt(x,t) = \alpha_{\rm w} - k_7\, sox(x,t) - k_9\, wnt(x,t) + D_w\, \partial_{xx} wnt(x,t). \tag{15}$$

We fix the parameters as in [16] of their Figure2b for which the systems exhibit a Turing instability. As there are no parameters which scale with the field-width, the length-scale of the pattern is independent of the field width. The number of digits would then be proportional to the width of the field, which is not what we observe. To this end we modify Eq. (13) such that the length-scale of the pattern scale with the field width, i.e. the number of digits is constant. A simple choice to achieve this behaviour is to include a scale factor $\gamma$ into the reaction terms [17],

$$\partial_t sox(x,t) = \gamma\big[\alpha_{\mathrm{s}} + k_2\, bmp(x,t) - k_3\, wnt(x,t) - \big(sox(x,t) - sox_0\big)^3\big] \tag{16}$$

$$\partial_t bmp(x,t) = \gamma\big[\alpha_{\mathrm{b}} - k_4\, sox(x,t) - k_5\, bmp(x,t)\big] + D_b\, \partial_{xx} bmp(x,t) \tag{17}$$

$$\partial_t wnt(x,t) = \gamma\big[\alpha_{\mathrm{w}} - k_7\, sox(x,t) - k_9\, wnt(x,t)\big] + D_w\, \partial_{xx} wnt(x,t). \tag{18}$$

In Fig. ST 7 we show that by choosing $\lambda \propto L^{-2}$ we retrieve perfect scaling of the pattern for increasing field width.

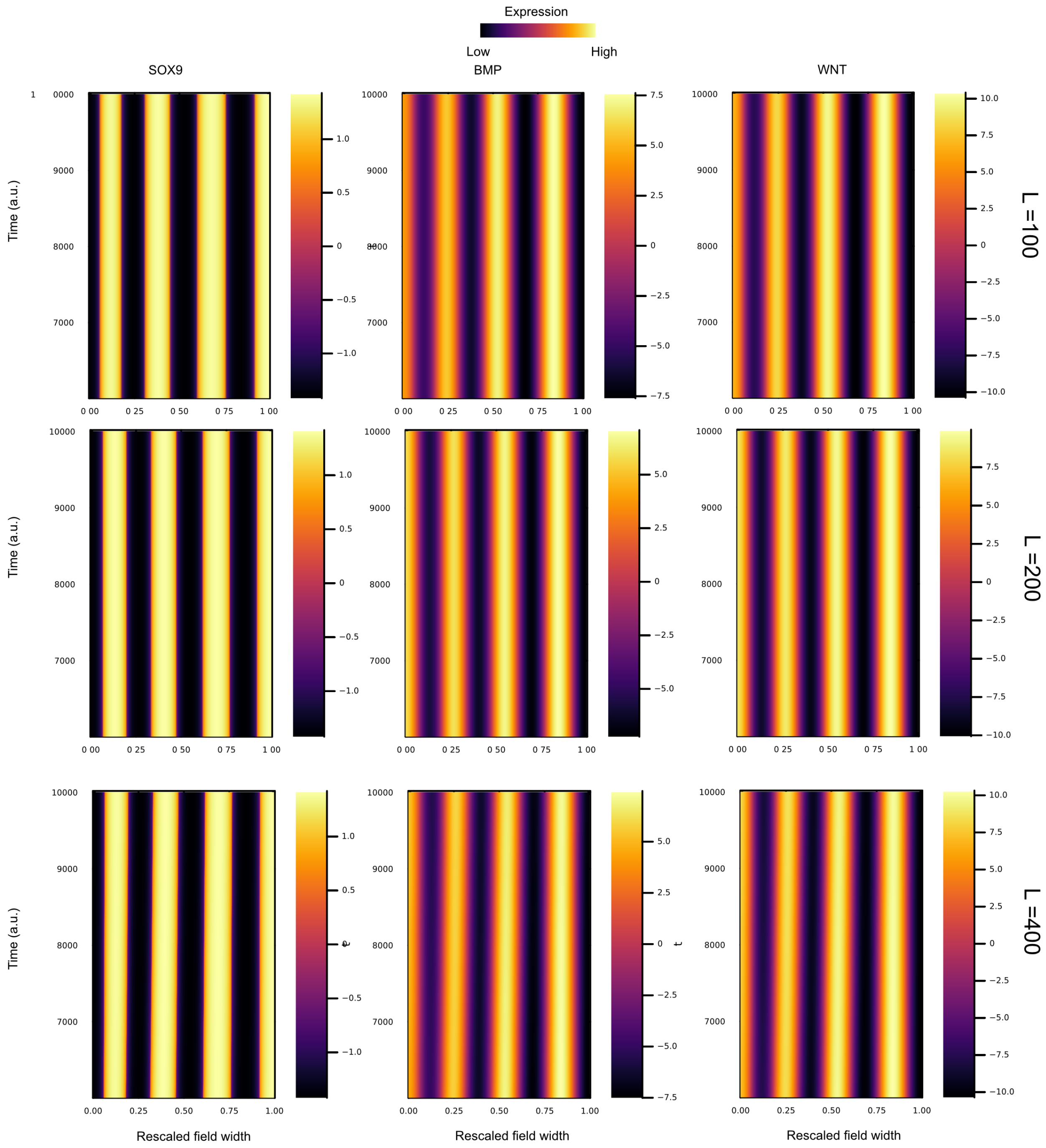


Figure ST 7: Simulations of Eq. (16) for three different system sizes, $L = 100, 200, 400$ reveals scaling of the digit period.

However, we found that larger animals have on average less digits than smaller animals. This seems to be in contraction with our previous simulations, but, as we shown in the main text, the field increase during digit specification is a function of the animal size. To this end, we need to include tissue growth into the Turing equations, as shown in Sec. 4.3.

#### 4.1 WNT perturbation

In this section we prove how a perturbation of WNT lead to diverging morphologies as observed experimentally (Fig. 3).

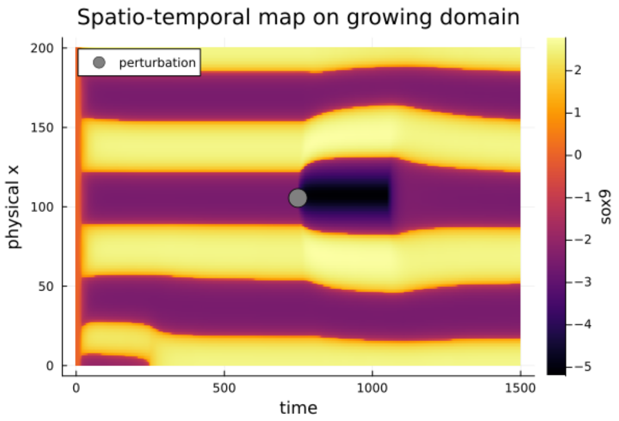


Figure ST 8: WNT inhibiton in the interdigit leads to an increase of the interdigit distance.

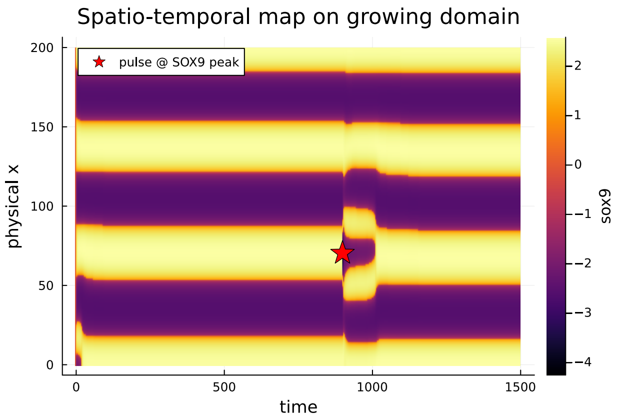


Figure ST 9: WNT inhibiton on the digitleads to a flattening of the domain, thus partially increasing the amplitude of the pattern.

In order to resemble as closely as possible the experimental perturbation, we modify the WNT equation as following,

$$\partial_t w(x,t) = \Big(\alpha_{\rm w} + \Delta\alpha_{\rm w}\, \Pi(t;t_0,T)\, g(x;x_0,\sigma)\Big) - k_7\, sox(x,t) - k_9\, w(x,t) + D_w\, \partial_{xx} w(x,t). \tag{19}$$

where

$$\Pi(t;t_0,T) = \begin{cases} 1, & t_0 \le t \le t_0 + T, \\ 0, & \text{otherwise}, \end{cases} \tag{20}$$

$$g(x;x_0,\sigma) = \exp\left(-\frac{(x-x_0)^2}{2\sigma^2}\right), \tag{21}$$

Specifically, we consider an injection ($\Pi$) of WNT at a given time and centered around one inter-finger zone $x_0$ and $\Delta\alpha_{\rm w}$ controls its strength. The results are shown in Fig. ST 8 where we observe a similar morphology as in the experiment (Fig3 in the main text). As the simulations are one-dimensional, time plays the role as the second spatial dimension. We then performed numerical simulations where the WNT injection is positioned on the finger rather than in the interdigit. This is done by simply shifting the value of $x_0$ in the previous equation. The resulting behavior is shown in Fig. ST 9 where again a short perturbation leads to finger deformation as observed experimentally in Fig. 3.

#### 4.2 Failure of scaling via SHH expander

Here we show that the loss of one digit and their scaling are not likely due or connected to the failure of scaling of the SHH expander. Specifically, if we couple the expander to the Turing system, a simple choice would be to have the scaling factor $\gamma \propto \bar{e}^{-2}$. This choice is made such that for a perfectly scaling expander, we recover Fig. ST 7. However, for an expander which growths sublinearly with the field width, $\bar{e} \propto L^{\alpha}$, $\alpha < 1$ the resulting pattern for the larger field width is the one shown in Fig. ST 10. As shown, in the case of a direct coupling to the expander, the larger animals would form more digits than less on average. We conclude that this simple choice is not suitable to explain the experimental observations.

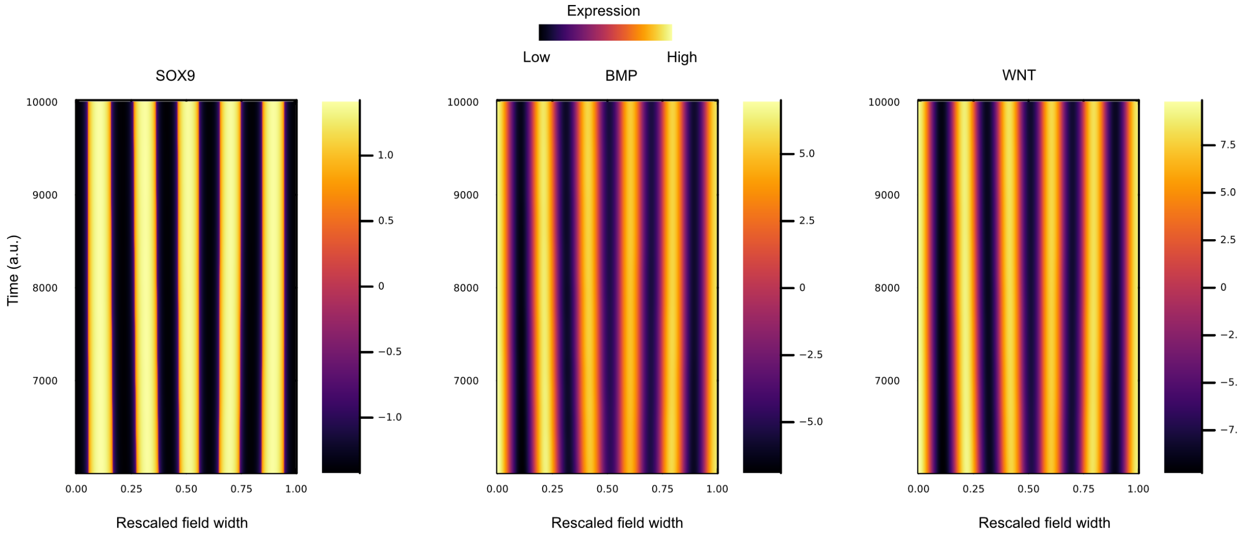


Figure ST 10: Coupling of Eq. (16) via a scale factor term proportional to the SHH expander, $\gamma \propto \bar{e}^{-2}$ reveals that larger animals are expected to have more digits.

#### 4.3 Turing models on growing domains

As explained in the main text, we found that the scaling of the digit period with a lack of a digit for the larger animals is due to insufficient growth of the blastema width. In order to model tissue growth, we need to consider that the length of the system $L$ is now a function of time $L(t)$. Mathematically, we can always solve on a domain $x \in [0, L(t)]$ by rescaling the domain into a time independent domain $\xi \in [0, 1]$ where $\xi = x/L(t)$. All the derivatives change accordingly $\partial_x^2 \to \partial_\xi^2/L(t)^2$. When passing from a fixed domain to a growing domain equations get an additional advective-like term coming from the comoving coordinate frame [18]. This term scales as $\partial_t L(t)/L(t)$. We note that there is a possible addition of other terms due to dilution, but this term does not come from coordinate transformation. In our case, the minimal tissue width measurement is approximately 1 mm and it growths by a factor of 1.4 in approximately 7 days. If we consider linear growth, this term is approximately $6 \cdot 10^{-7} s^{-1}$. If we consider a very slowly diffusion protein (diffusion of order $10^{-12}$) than $D/L^2 \approx \cdot 10^{-5} s^{-2}$, so it is approximately fifty times larger. We can then safely neglect the advective-like term for digit patterning during axolotl limb regeneration.

Moreover, we have to take into account that the pattern is scaled such that the scale factor $\lambda$ are proportional to the inverse squared system size as we shown before, i.e. $\lambda \propto L^{-2}$. Here lies the more subtle part, as we understood that despite the system is able to scale proportionally to the initial field size, a failure of growth impact the scaling. To this end, we fix the growth to be $\lambda \propto L_0^2$. In this way we recover early scaling, Fig. ST 11a. On the other hand, if we would consider the unphyiscal limit of a tissue expanding 10 fold than more digits will appear,Fig. ST 11a. This is opposed to the case where $\lambda \propto L(t)^{-2}$ where the pattern will scale indefinitely, Fig. ST 11b

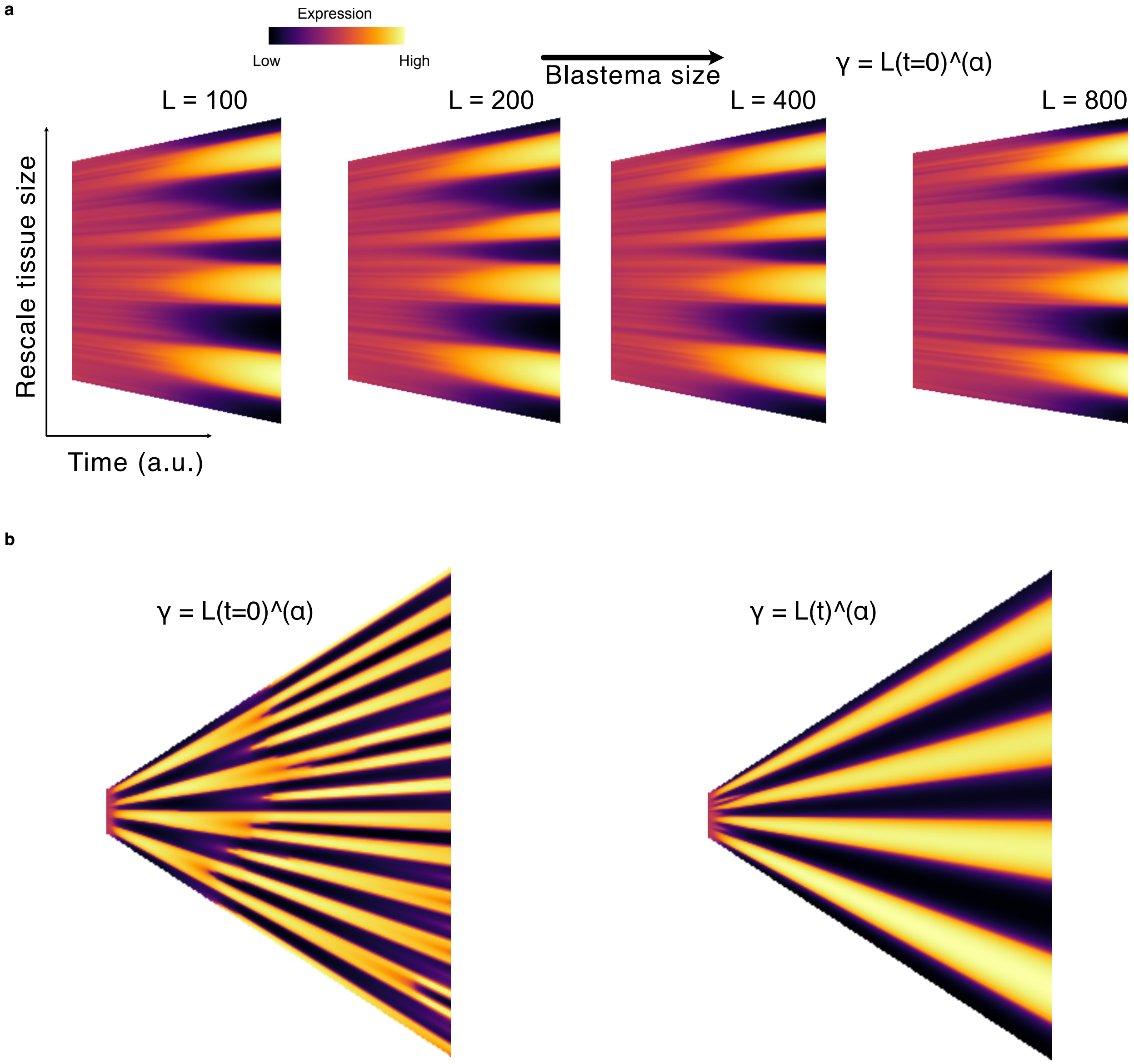


Figure ST 11: **a** Scaling of Turing pattern with tissue growth for an eight-fold-change of tissue sizes. The growth is stopped when the final tissue size is 1.4 greater than the initial as measured experimentally. We set growth rates to scale accordingly in order to recover almost perfect scaling between tissue sizes. **b** For the nonphysical scenario of tissue growing beyond the limits, if $\lambda \propto L(t)^{-2}$ we would observe a perfect scaling. On the other hand, with our choice of $\lambda \propto L(t=0)^{-2}$, the pattern bifurcates.

#### 4.4 Sequential digit appearance

Turing patterns are rarely reproducible exactly if initial conditions are taken to be random perturbation over a stationary state. Moreover, the appearance of the peaks of the pattern is generally simultaneous. However, we observed that the digit sequence appear in the order 2-1-3-4 where the fourth digit is close to the SHH domain. In order to recover the right digit sequence we ought to take the minimal modelling choice, which is to have a 4-th gene domain (HOX as suggested in [16]) which initially is placed around the zone of the 2nd digit and then expand till almost reaching the boundary of the domain. To do so we introduce a dynamical equation of the HOX front. Parameters are changed accordingly as in [16] and the final results are shown in Fig. ST 12a when there is tissue growth and in Fig. ST 12b, without tissue growth.

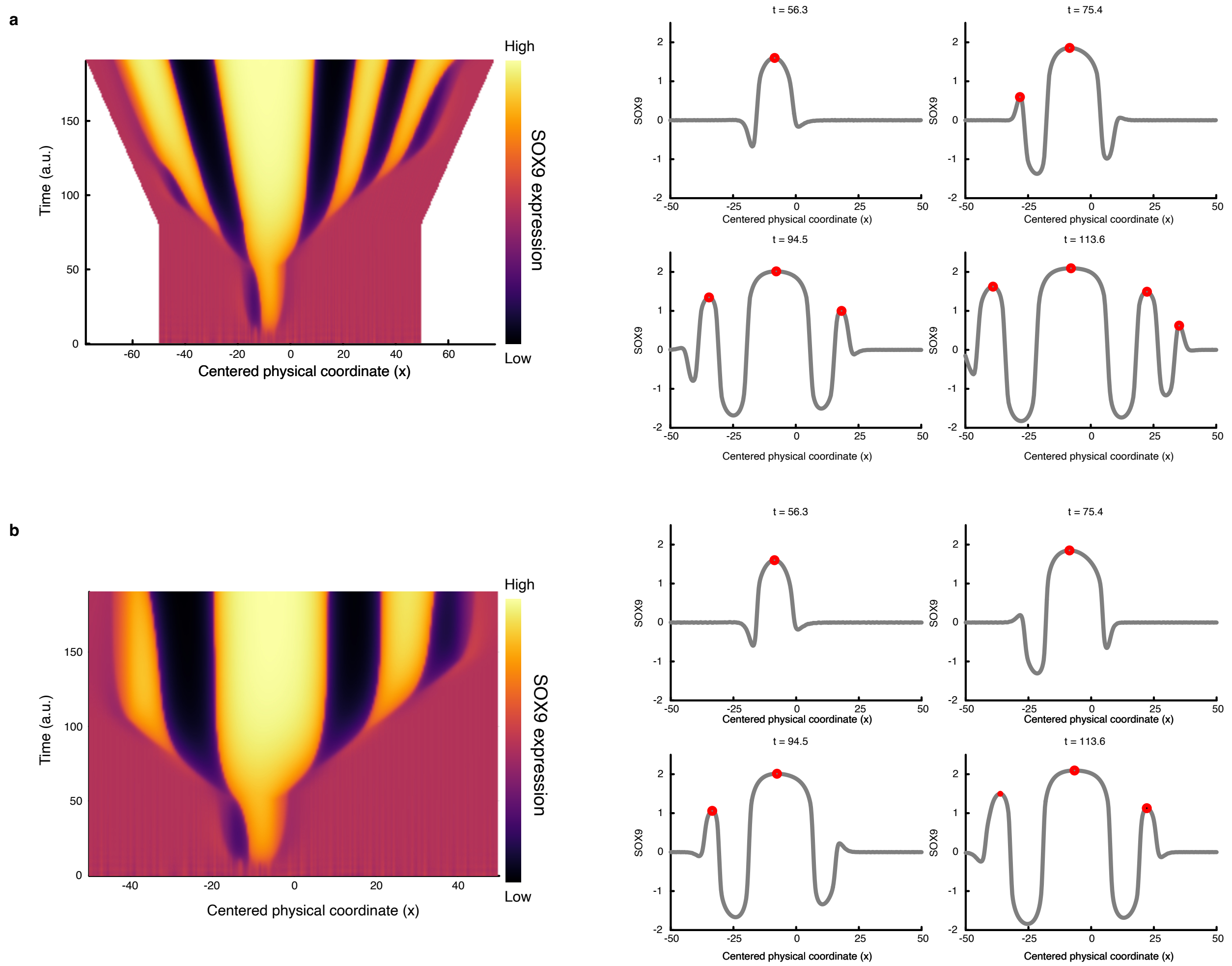


Figure ST 12: **a** HOX domains couples to tissue growth and produce sequential digit appearance 2-1-3-4. **b** Digits appears sequentially even without tissue growth and a lack of the 4th digit.

Here we do not claim that this is the only mechanisms giving rise to sequential digit patterning and other mechanisms including possible couplings to the SHH and FGF pathway might lead to the initial instability zone.

##### 4.5 Alternative network architectures

Here we used the minimal topology proposed in [16] which gives rise to patterning of the digit and we introduce the effect of scaling of parameters as well as growth. We argue that despite that architecture give rise to the observed patterns, more complicated architectures for example involving INHBA instead of BMP or both genes could give rise to the exact same phenomenology. Theoretically, how network topology selects different in-phase and out-of phase patterns (as shown in Fig. S4d of the main text) was studied in [19]. As our aim was to show that such models could give rise to scaling of the digit period and lack of a digit in case of a failure of growth, we do not claim that the precise network architecture chosen is the most realist one for digit patterning during axolotl limb regeneration. However, such architecture is minimal enough to correctly account the observed measurements.
